# Physiological and transcriptomic variability indicative of differences in key functions within a single coral colony

**DOI:** 10.1101/2021.02.25.432919

**Authors:** Jeana L. Drake, Assaf Malik, Yotam Popovits, Oshra Yosef, Eli Shemesh, Jarosław Stolarski, Dan Tchernov, Daniel Sher, Tali Mass

## Abstract

Polyps in different locations on individual stony coral colonies experience variation in numerous environmental conditions including flow and light, potentially leading to transcriptional and physiological differences across the colony. Here, we describe high-resolution physiological measurements and differential gene expression from multiple locations within a single colony of *Stylophora pistillata*, aiming to relate these to environmental gradients across the coral colony. We observed broad transcriptional responses in both the host and photosymbiont in response to height above the substrate, cardinal direction, and, most strongly, location along the branch axis. Specifically, several key physiological processes in the host appear more active toward branch tips including several metabolic pathways, toxin production for prey capture or defense, and biomolecular mechanisms of biomineralization. Further, the increase in gene expression related to these processes toward branch tips is conserved between *S. pistillata* and *Acropora* spp. The photosymbiont appears to respond transcriptionally to relative light intensity along the branch and due to cardinal direction. These differential responses were observed across the colony despite its genetic homogeneity and likely inter-polyp communication. While not a classical division of labor, each part of the colony appears to have distinct functional roles related to polyps’ differential exposure to environmental conditions.

## 1 Introduction

Coral reefs remain one of the most biodiverse ecosystems on Earth (Knowlton et al., 2010), despite facing a suite of anthropogenic stressors with both local and global causes (e.g., Hughes, 1994;Hoegh-Guldberg et al., 2019). Most stony corals (order Scleractinia), the sessile builders of the reef system, grow as colonies of thousands of genetically identical polyps formed by budding, with colonies ranging from centimeters to meters in diameter. The reef-building corals are also host to intracellular symbiotic dinoflagellates whose photosynthetic activity restricts the coral holobiont to the top ~100 m of the ocean and fulfills much of the host’s carbon nutritional requirements (e.g., Falkowski et al.). Nevertheless, active feeding on particles from the water column plays a significant role as well (e.g., Martinez et al.), with active predation utilizing toxins produced and delivered by cnidocytes (recently reviewed by Schmidt et al.). These fixed carbon sources ultimately power the production of the external calcium carbonate skeleton that forms the reef and persists after the animal has died.

While the individual polyps in most colonial corals are genetically identical, each polyp inhabits a distinct location within the colony, which can be morphologically complex (e.g., in the case of branching corals). Thus, it has been suggested that individual polyps within a coral colony are differentially exposed to environmental conditions such as water flow and light availability (e.g., Carpenter and Patterson, 2007;Chang et al., 2009), which have distinct effects on coral physiology. These micro-environmental conditions may play a role in feeding capability (Sebens et al., 1997), reproductive success (Mass et al., 2011), photosynthesis (Dennison and Barnes;Mass et al., 2007), calcification rate (Dennison and Barnes, 1988), and waste removal (Mass et al., 2010). For instance, reductions in flow between the colony branches can trap food particles, making it easier for hunting tentacles of the polyps to efficiently pull them from the water column (Chang et al., 2009), and leading to downstream effects on metabolism (e.g., Gladfelter et al., 1989). Similarly, differences in light exposure due to the coral’s branching leads to self-shading that can dramatically reduce light availability (Anthony et al., 2005), potentially leading to fine-scale spatial heterogeneity in photosynthetic parameters between individual polyps as well as between polyps and the coenosarc (Ralph et al., 2002). We hypothesize that such physiological differences will be underpinned by transcriptional differences. Plasticity in host gene expression has been observed at coral branch tips versus bases across multiple colonies (Hemond et al., 2014). However, it has not previously been linked, at high spatial resolution, to physiological and morphological measurements indicative of relevant environmental parameters and physiology within a single coral colony.

Here we combined high-resolution transcriptomic analysis with field measurements and physiological assays to describe intra-colony plasticity in the Indo-Pacific stony coral, *Stylophora pistillata*. We sought to answer two questions: first, are there differences in function by groups of polyps at different locations throughout the colony; and secondly, can we attribute those differences to specific location characteristics that are likely associated with varying microenvironments across the colony? Our whole-genome expression and physiological data are from a single coral colony at three locations along each branch taken from a different location within the colony. This high-resolution dataset enables us to resolve variability along three main axes – tip-junction-base, top to bottom, and the cardinal direction (Figure 1). Each of these axes is associated with changes in gene expression and physiology or morphology, which we suggest are due in part to light and current direction and strength. It appears that polyps in the interior of the colony are exposed to relatively stagnant conditions, whereas the colony periphery likely experiences higher flow, light exposure, chances for predation, and requirements for competition and defense leading to the dynamic physiological and transcriptional responses observed here. Our findings have implications for the interpretation of data from branches collected from different locations across multiple coral colonies. Specifically, when fragments are taken from multiple colonies for replication, considerations of the inherent differences across a colony should be included in sampling design. Our findings also shed new light on the potential drivers of plasticity of corals’ responses to environmental stress and disease, and should be considered when reconstructing paleophysiology and past microenvironmental conditions from well preserved fossil coral colonies.

**Figure 1.**
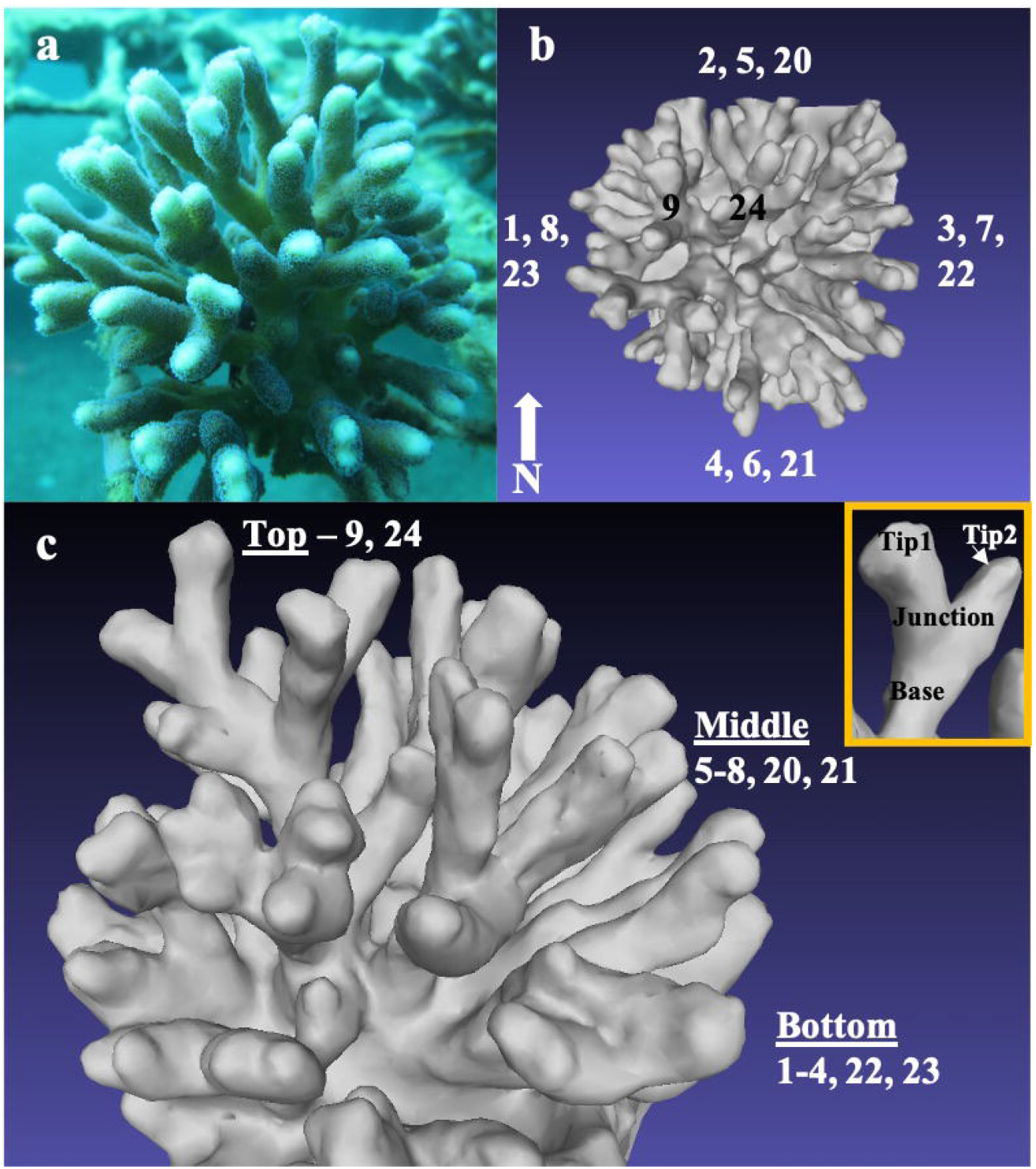
Micro-environments and responses in sections of an individual coral colony. Photograph of the *S. pistillata* colony analyzed for physiological and transcriptomic intracolonial differences taken *in situ* (8 m depth) from the east side of the colony (a). Three-dimensional reconstructions of the coral colony showing locations of branches based on cardinal direction (top view, b) and ring height (oblique view, c). Inset in (c) shows branch axis sampling locations at tips, junction, and base per branch.

## 2 Manuscript Type

Original Research, Body text is 8048 words.

## 3 Methods

### 3.1 Field measurements

#### 3.1.1 Sample collection

Ex situ measurements were conducted on a 15 cm diameter colony of *Stylophora pistillata* collected during the morning in April 2016 at 8 m depth under a special permit from the Israel Nature and Parks Authority in the coral nursery of the Inter-University Institute for Marine Science (IUI), Eilat, Israel using SCUBA (Figure 1a). The colony was brought to the lab in a tank filled with seawater and divided into individual branches and branch segments within 20 minutes of collection. Nine branches (numbered 1-9 in Figure 1b, c and supplementary materials) were taken from three horizontal concentric rings (one ‘top’ branch, four ‘middle’ branches, and four ‘bottom’ branches) and classified according to their relative height within the coral colony (Figure 1b, c). The eight middle and bottom branches also represent duplicates from each cardinal direction (N, S, E, W). All branches were further divided into sections according to their position on the branch (tip, junction, base) and flash-frozen in TRI-Reagent (Sigma) (Figure 1 inset) for differential gene expression analysis. Five additional branches (numbered 20-24 in Figure 1b, c and supplementary materials) distributed around the colony were chosen for physiological examination, fragmented into tip, junction, and base segments, and then kept at – 80 °C.

#### 3.1.2 Oxygen measurements

Immediately prior to removal of the colony intended for high spatial resolution examination from the coral nursery, dissolved oxygen (DO) in seawater between branches of five *S. pistillata* colonies located in the IUI coral nursery was measured in-situ. This nursery is a pair of raised 4×3 m frame structures (part of one can be seen behind the coral colony in Figure 1a) ranging in depth from 5-10 m, so that conditions experienced by all well-spaced colonies are quite similar across the nursery. Measurements were taken as up to five radial transects per colony, from inside to just outside each colony (i.e., center, mid-way, and periphery), using Unisense oxygen microsensors mounted on a Unisense UnderWater Meter System. The microsensors were moved through each colony at a controlled pace in order to keep the intra-colony water as undisturbed as possible and to prevent exchange of inter-branch water within the colony and with the surrounding water. Each measurement was taken for a duration of two minutes to allow the oxygen microsensor readings to stabilize. All measurements for each transect were normalized to the outermost reading for that transect (Supplementary Table 1). DO concentrations across one representative transect were calculated according to the following equation to establish that such values were normoxic: DO (μmol/L) = (DO_sat_(μmol/L)/(mV_sat_-mVo)) * in situ mV measurement.

#### 3.1.3 Microbial counts

Microbial counts outside and within the coral colony were performed at the same time as DO measurements on samples collected in-situ via radial transects on five different colonies growing at the same site. We conducted preliminary experiments using small aliquots of concentrated fluorescein dye injected to water between branches of *S. pistillata* skeletons, the same size as that used in our physiological experiments, in a static laboratory aquarium to test the efficacy of extracting water from between branches and at colony peripheries with minimal intra-colonial water mixing. Briefly, we injected the dye on one side of a branch and then used a syringe to carefully extract various volumes of water, from 0.4 to 12 ml in intervals of 0.4 ml, from the other side of the branch, with five replicate experiments per extract volume tested. We then measured the fluorescence of the extracted water, subtracted the fluorescence background of the surrounding water, and represented the results as a percentage of the fluorescence from the injected fluorescein dye aliquot. This showed that we could extract up to 1.6 ml of seawater from between coral branches and from colony peripheries without pulling in water from the immediate environment of the surrounding branches (Supplementary Figure 1). Water flow during our sampling dive was noticeably very low. We therefore felt confident extracting 1 ml of seawater from between the branches (‘center’ position) and the tip of the branch at the colony periphery (‘periphery’ position).of the live colonies. Upon completion of the dive, all water samples were immediately carried to the lab and fixed in cryovials in a final concentration of 0.0625% glutaraldehyde, incubated in the dark for 10 minutes, and then flash-frozen at −80 °C until analysis by flow cytometry.

Just prior to analysis, microbial count samples were thawed at room temperature and diluted 1:1 in ultra-pure water, and 2 μm-diameter fluorescent beads were added to each sample as an internal standard (Polysciences, Warminster, PA, USA). The samples were analyzed on a FACSCanto™ II Flow Cytometry Analyzer Systems (BD Biosciences). We first examined the natural fluorescence of the cells (chlorophyll and phycoerythrin pigments). We then stained the cells with SYBR Green I (Molecular Probes/ThermoFisher) according to the manufacturer’s instructions and counted the total microbial population as well as cyanobacterial sub-populations. Data were acquired and processed with FlowJo software. Flow rates of the instrument were determined several times during each run and the average value for a sample-free test run was used for calculating cell per ml.

### 3.2 Physiological analysis

#### 3.2.1 Tissue removal

Tissue was removed from five frozen coral fragment skeleton by airbrush with phosphate buffered saline (PBS) and homogenized using an electrical homogenizer (MRC, HOG-160-1/2) for 10 seconds. A sub-sample of each fragment’s homogenate was stored for cell counts, chlorophyll extraction, and total protein measurements and the rest of the homogenate was centrifuged for 5 minutes at 5000 rpm. The supernatant was retained for host protein and hemolysis assay. The pellet was observed under the microscope to quantify photosynthetic symbionts. All quantities were adjusted to fragment extraction volume and normalized to cm^2^ of the fragment’s surface area.

#### 3.2.2 Chlorophyll extraction

Two ml of tissue homogenate was filtered on a Whatman GF/C filter and incubated with 1 ml 90% acetone for two hours at 4°C. After incubation, the filter was manually homogenized and the solution was filtered through a 0.22 μm syringe filter into a glass cuvette. Spectrophotometric measurements were conducted on a NanoDrop (Thermo-Fisher) and chlorophyll a concentrations were calculated from light absorbance results based on the following equation (Ritchie, 2008): chl-a [mg/ml]= −0.3319(ABS630)-1.7485(ABS_647nm_)+11.9442(ÄBS_664nm_)-1.4306(ABS_691nm_).

#### 3.2.3 Nematocysts and photosynthetic symbiont counts

100 μl of tissue homogenate was used for nematocyst (supernatant) and photosymbiont (pellet) counts by hemocytometer (BOECO, Germany); cells were counted in five randomly selected fields per fragment (1 mm^2^ each) on a Nikon Eclipse Ti-S Inverted Microscope System. Nematocysts were counted under white light with a contrast filter whereas photosynthetic symbionts were viewed by fluorescence at 440 nm excitation and 590 nm emission. NIS ELEMENTS (Nikon) software was used for the cell counts with automatic counting settings limited to cell diameters smaller than 15 μm and circularity set to >0.5 but <1.

#### 3.2.4 Total protein and host protein

We quantified homogenate and host total protein; sonication (Ultrasonic Atomizer Probe Sonicator) was used for further extraction of symbiont protein. Protein concentration of each sample was measured by bicinchoninic acid (BCA) assay (Pierce) against a bovine serum albumin standard curve at an absorbance of 540 nm.

#### 3.2.4 Hemolysis assay

Hemolytic assays were performed against O Rh positive human blood cells obtained from the Rambam/Yoseftal Hospital Blood Bank, as described previously (Primor and Zlotkin, 1975). Briefly, 2 ml of whole blood was diluted to a final volume of 15 ml in PBS (pH 7.4) and centrifuged at 3000 g for 5 min. The supernatant was removed, and the process was repeated with the pelleted erythrocytes until the supernatant was clear. Washed erythrocytes were then resuspended in PBS to a final concentration of 20% (v/v). For each hemolysis assay, 160 μl of homogenate supernatant was incubated with 40 μl of washed erythrocyte suspension (4% erythrocytes v/v) at 37°C for 30 minutes in a water bath. At the end of the incubation, 400 μl of PBS was added, and the assays were centrifuged at 3000 g for 3 min. The supernatant fluid containing the hemoglobin released from lysed erythrocytes was transferred to 96-well microplates and the absorbance at 540 nm was determined on a spectrophotometric microplate reader (Perkin-Elmer). In addition, each experiment was normalized to a positive (100% hemolysis) and negative control (0% hemolysis) by incubating erythrocytes with DDW and PBS alone, respectively. HU50 was defined as the amount of homogenate required to cause 50% hemolysis (Bartosz et al., 2008) in a dilution series of the coral extract.

#### 3.2.5 Fragment surface area

Surface areas were estimated using the aluminum foil method (Marsh Jr, 1970). Briefly, aluminum foil was molded over the skeleton of each fragment (without measuring skeleton connective areas between the divided fragments), carefully removed, and weighed. This was repeated three times on every fragment and the surface area was estimated using a standard curve of the derived relationship between foil area and weight.

#### 3.2.6 Skeleton cleaning for assessments of surface area and corallite measurements

After tissue extraction, all fragments were cleaned of organic residues overnight in 3% sodium hypochlorite; the skeletons were then washed in deionized water and dried at 55°C.

#### 3.2.7 Micromorphological analysis

The distance between neighboring polyps and the surface area of the polyp was measured using a Nikon binocular microscope and analyzed using NIS-Elements (Nikon) software. For each coral fragment, at least four corallites located in a position parallel to the focal plane were chosen. Major and minor axes and circularity were measured for each corallite and the ratio of the two axes (major divided by minor) was calculated; a ratio of 1 indicates a circle and the larger the ratio, the more elliptical the shape. Surface area of each measured corallite was calculated from the two-measured axes, using the following equation: S = π* r (major axes)* r (minor axes). Finally, distances between neighboring corallites were also measured to evaluate polyp density in each area. Corallites were qualitatively analyzed by scanning electron microscopy.

### 3.3 Differential Gene Expression Analysis

#### 3.3.1 RNA extraction, processing, and sequencing

Nine branches, each segmented into three portions along their axes, yielded 36 transcriptome libraries across the three-dimensional space of the coral colony (Figure 1, colony locations in Supplementary Table 3, read count statistics in Supplementary Table 4). Total RNA was extracted from the holobiont in fragments stored in TRI-Reagent (Sigma) following the manufacturer’s protocol, with some modification at the homogenization step. Briefly, samples frozen in TRI-Reagent were heated and centrifuged, and then bromochloropropane, at a ratio of 1:10, was added to the samples for separation. After incubation at room temperature for 10 minutes and centrifugation, RNA was purified from the clear phase using a Purelink RNA Mini Kit (Ambion) according to the manufacturer’s protocol. The RNA was washed in 70% ethanol and then on-column DNase digestion was performed using a Qiagen RNase-free DNase Kit according to the manufacturer’s protocol. RNA-Seq libraries were prepared using an in-house protocol at the Weizmann Institute of Science. Briefly, the polyA fraction (mRNA) was purified from 500 ng of total RNA per sample followed by fragmentation and generation of double-stranded cDNA. Then, end repair, A base addition, adapter ligation and PCR amplification steps were performed. Libraries were evaluated by Qubit (Thermo Fisher) and TapeStation (Agilent). Sequencing libraries were constructed with barcodes to allow multiplexing of all samples to be run in each lane. Approximately 578 million total high-quality 125 bp paired end reads (15.45 ± 1.5 million paired reads per sample) were sequenced on an Illumina HiSeq 2500 across three different lanes (i.e., each sample run in triplicate to remove batch effects).

#### 3.3.2 Host and symbiont RNA-Seq differential expression analysis

Standard RNA-Seq quality filtering was conducted as described previously (Malik et al., 2021). Briefly, RNA-Seq reads were adapter-trimmed using cutadapt 1.15 (https://cutadapt.readthedocs.io), and then low-quality regions were removed with Trimmomatic 0.3 (Love et al., 2014). In order to detect correct taxonomic classifications within our transcriptome data, high-quality reads were mapped to all available NCBI and Reefgenomics (reefgenomics.org; (Liew et al., 2016)) genome-based proteome databases of Symbiodiniaceae species *Symbiodinium microadraticum, Cladocopium goreaui*, and *Fugacium kawagutii* (formerly *Symbiodinium* spp. clades A, C1, and F, respectively (LaJeunesse et al., 2018)), and Cnidaria as well as selected stramenopiles/alveolates/Rhizaria and Metazoa databases using Diamond (Buchfink et al., 2015). Top hits almost exclusively belonged to robust corals (mainly *S. pistillata*), and *S. microadriaticum* (formerly “clade A” (LaJeunesse et al., 2018)). These results are expected since *S. microadriaticum* was found to be a predominant symbiont in shallow water in Eilat (Malik et al., 2021). For symbiont RNA-Seq mapping, we further aligned the reads to the merged host genome assembly (NCBI GCA_002571385.1) and the symbiont genome assembly (NCBI GCA_001939145.1) using STAR (Dobin et al., 2013;Aranda et al., 2016;Voolstra et al., 2017). Across all branch fragments, 72-80% (~11-16 million reads) were concordantly mapped to the host genome, and 2%-10% (~0.5-1.5 million reads) were concordantly mapped to the symbiont genome. In total, 83-84% of the reads were aligned to either host or symbiont genomes. Differential expression (DE) analysis was conducted using Bioconductor DEseq2 (Love et al., 2014), separately for the host and the symbiont genes, using a DEseq2 generalized linear model. Specifically, we tested: (1) the additive effect of branch position, ring height and cardinal direction; (2) the additive and interaction effect of branch position and ring height (excluding cardinal direction factor); (Supplementary Tables 5-7). Note that the first model cannot include the four top branch samples (two tips, junction, base), since the single top branch is represented by an incomparable cardinal direction (“center”); as such, these samples were excluded from the analysis of expression of putative toxins and known biomineralization-related genes. For the symbiont analysis, only the first model was used. Although DESeq2 is designed to analyze samples of variable total read counts, in order to avoid read normalization problems, we only selected symbiont genes whose 25% quantile of read count was at least 12 reads (with 12,735 symbiont genes remaining); further, only symbiont samples with >300k total reads were retained. After filtering, the correct symbiont read normalization was indicated by a typical close-to-symmetric distribution of fold-changes around the DESeq2 MA plot horizontal axis (Supplementary Figure 2). NMDS analysis was conducted using metaDMS in the R Vegan package based on log_10_FPM values of all expressed genes. For the symbiont NMDS, since read counts were relatively low, pairs of tips from the same branch are represented together based on the sum of read counts for both tips; however, symbiont DESeq2 data were not pooled for differential gene expression analysis.

#### 3.3.3 Branch genetics and SNPs analysis

Single nucleotide polymorphisms (SNPs) analysis was conducted for each sample using the Broad Institute-recommended RNA-Seq SNPs practice (https://gatk.broadinstitute.org). The pipeline includes four steps: STAR reads mapping, a pre-processing step aimed at removing various alignment biases, variant calling using GATK version 3.5, and finally variant filtration and annotation (DePristo et al., 2011). In order to exclude variant-call biases due to changes in transcript abundance between samples, we considered only SNP loci with coverage greater than 30 RNA-Seq reads in all tested samples. Identity By Decent (IBD) analysis was conducted using SNPRealte in R.

#### 3.3.4 Host and symbiont functional enrichment analysis

Biological terms were assigned to genes based on Uniprot *S. pistillata* (www.uniprot.org), KEGG *S. pistillata* (www.kegg.jp), *S. pistillata* Trinotate annotations (Bryant et al., 2017), and Uniprot *S. microadriaticum* databases. Enrichment analysis was conducted in Bioconductor GOSeq (Young et al., 2012), which corrects for enrichment biases associated with correlations between gene size and DE significance. We also searched for functional enrichment using the score-based tool GSEA (Subramanian et al., 2005;Sergushichev, 2016), which may be more sensitive than p-value cutoff-based searches (such as GOSeq) when small non-significant changes in relatively large groups of genes are expected. In GSEA, we used log2 fold changes as scores. Because many enriched terms are functionally related, we hierarchically clustered the biological terms based on pairwise distances between groups of genes: D = |xa⋂xb|/minimum(|a|,|b|), where a,b are two sets of gene ids, and xa and xa are DE genes ids from a and b, respectively. Terms trees were constructed using Bioconductor-ggtree (Yu et al., 2018). Terms enriched in ring height comparisons to the top of the colony should be viewed with caution as only one branch was taken from that classification. Only significant enrichment results are reported.

#### 3.3.5 *Focused* S. pistillata *gene analyses*

Putative toxin genes were identified using reciprocal best-BLAST hits as described in (Rachamim et al., 2015;Yosef et al., 2020) (Supplementary Table 14).Differential expression of genes coding for known biomineralization-related proteins from *S. pistillata* skeleton or the calicoblastic layer were examined (Puverel et al., 2005;Drake et al., 2013;Mass et al., 2013;Zoccola et al., 2015;Peled et al., 2020) (Supplementary Table 16).

### 3.4 Statistical Analyses

Statistical tests of all skeletal, physiological, and environmental data described above were performed in RStudio (RStudio Team). If a parameter displayed both homogeneity of variance (Fligner-Killeen test) and normal distribution (Shapiro-Wilke test), a one-way ANOVA (chlorophyll/total protein, chlorophyll/photosymbiont cell, photosymbiont/surface area, total protein/surface area, host protein/surface area, nematocysts/total protein, nematocysts/surface area, corallite diameter, corallite distanc) was performed, with TukeyHSD post hoc analysis for parameters with significant differences between locations. Parameters with non-homogenous variance and/or non-normal distribution were analyzed using Wilcoxon’s rank sum test (water column microbes, DO) or the Kruskal-Wallis rank sum test (chlorophyll/surface area, photosymbiont/total protein, hemolytic units/total protein, hemolytic units/surface area, corallite axis ratio), with Dunn’s test for post hoc analysis for parameters with significant differences between locations. Environmental parameters were pooled between all colonies and tested between measured locations as described above. Coral and photosymbiont physiological and morphological parameters were tested between branch axis locations across the colony (i.e., five replicates each of junctions and bases, and 10 replicates of tips); however, there was not sufficient replication at cardinal directions or ring height for statistical evaluation of parameters between these locational classifications.

## 4 Results and Discussion

### 4.1 Differences in water DO and microbial counts across colony structure

Because branching corals’ three-dimensional structure can lead to different micro-environments within the colony, we first aimed to characterize indicators of the environmental conditions experienced by polyps at different locations within the colony that could potentially affect their gene expression. Rather than making direct measurements of flow within *S. pistillata* colonies at the IUI reef, we instead measured several within-colony water parameters affected by flow. We focused on dissolved oxygen because its concentration can vary within coral colonies, affecting coral physiology (e.g., Mass et al., 2010). Likewise, microbial counts represent an easy to quantify variable that can be used as a proxy of water stability. We measured higher oxygen concentrations in water collected close to the core (center; ~250 μmol/L) of the corals, compared to samples collected at colony peripheries (~230 μmol/L; p<0.05, Wilcoxon rank sum test; Figure 2a; Supplementary Tables 1, 2). Although there was variability in DO signal between colonies (Supplementary Figure 3), the ‘center’ versus ‘periphery’ differences held across all colonies. We also observed higher microbe densities in water collected from the colony interiors versus the immediately surrounding seawater (p<0.05, Wilcoxon rank sum test; Figure 2b; Supplementary Tables 1, 2). Both of these observations may be due to lower rates of water exchange at branch bases due to decreased water flow within the colony, as suggested by (Chang et al., 2009). Lower mass transport has been observed within the less densely branched *Pocillopora eydouxi* whose overall colony structure is similar to the *S. pistillata* studied here (Figure 1a) compared to the more densely branched *P. meandrina* (Hossain and Staples). Flow reduction within the colony would cause the accumulation of oxygen produced by photosynthesis (Mass et al.) and we speculate might also support the retention of microbes actively growing between branches (e.g., Schiller and Herndl;Ritchie, 2006), or those chemotaxing toward the coral tissue (Garren et al., 2014). These differences further support the notion that the environment differs along the coral branches from colony exterior to interior and may affect the coral holobiont with some polyps inhabiting more or less favorable areas, which we observe being reflected in transcriptomic and physiological differences across the colony.

**Figure 2.**
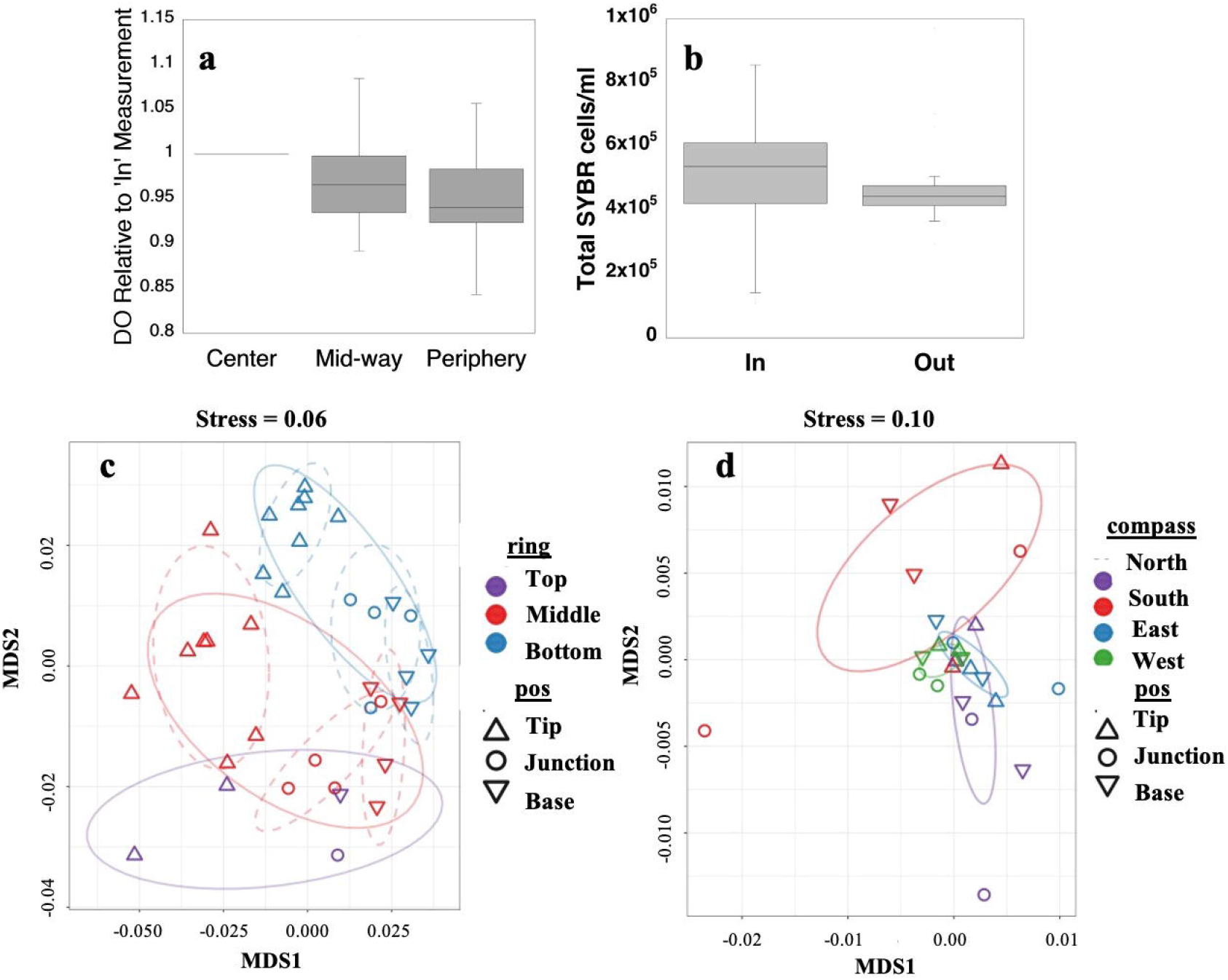
In-situ water column concentrations of dissolved oxygen (a) and total microbial cells (b) between branches inside and outside *S. pistillata* colonies. Non-metric multidimensional scaling (NMDS) of all expressed genes of the coral host (c) and photosymbiont (d) with factor effects color- and shape-coded. Host ring and branch position-based clusters are significant (Supplementary Table 7) and marked by dashed and solid ellipses, respectively.

### 4.2 Location-specific intracolonial variability of physiology and gene expression

To assess how the differences in environmental conditions resulted in different physiology and gene expression across a colony, immediately after measuring DO and collecting microbial samples we removed a single *S. pistillata* colony from the IUI reef, We then examined its branches both at different cardinal directions and heights above the substrate (i.e., ring height), with each branch further segmented along its axis from base to tip (Figure 1 inset; Supplementary Table 3). The identified patterns in gene expression and physiology therefore represent responses to microenvironments experienced by the colony at the specific time of collection and depth at which the colony was growing. The *S. pistillata* colony examined here was genetically homogenous (not chimeric, Supplementary Figure 4), and thus any changes in physiology or gene expression between regions were likely a response of the polyps to their local micro-environmental conditions rather than genotypic differences. While our study was limited to a single colony to achieve high spatial resolution of data, many similar genetic patterns are conserved between two major stony coral clades (see below). Additionally, differences were observed in hemolytic potential between branch tips and bases, which were seen also in two additional colonies from an independent study (Ben-Ari et al.). Taken together, it is clear that there are transcriptional and physiological differences between various locations within the coral colony, and that many of these differences are likely to be common across branching colonial corals. The expression of >10,000 of the ~24,000 tested host genes was affected by at least one of the tested factors: location at branch, ring height, and cardinal direction (Supplementary Tables 4-6). We observed a clear differentiation in host gene expression between branch positions and from the top to the bottom of the colony (Figure 2c, Figure 3, Supplementary Table 5), although comparisons to the lone top branch should be viewed with caution. Combined, we observed an interaction effect between factors (Supplementary Tables 4, 6), as, for instance, groups of genes involved in signaling pathways important in development that were significantly enriched in branch tips versus bases as compared to middle versus bottom interactions (Figure 3b, Supplementary Figure 5). Comparing colony interior with periphery, host gene expression patterns are more similar between junction and base branch positions compared to tips, although not the same at junctions and bases (Figure 2c). In contrast to the roughly ½ of all tested genes showing differential expression between location classifications, the only physiological parameters that differed along branch axis were hemolytic potential and skeleton morphology (described in detail below). Elucidating the microenvironmental drivers of these genetic and physiological differences across a colony may aid in, for example, understanding spatial differences of polyps’ interactions with various infectious organisms that target the oral ectoderm (e.g., Gibbin et al.) leading to diseases such as skeletal band eroding disease (Winkler et al., 2004), which has been observed to commence at colony interiors. Other previously reported response differences across colonies that may be due to microenvironmental variations include bleaching that commences at colony perimeters in *Oculina* (Shenkar et al., 2005), greater storage of lipids and total fatty acids toward colony interiors in *Acropora* (Conlan et al., 2018), and differing relative abundance of various photosymbiont species in regions that later displayed differential bleaching susceptibility (Kemp et al., 2014).

**Figure 3.**
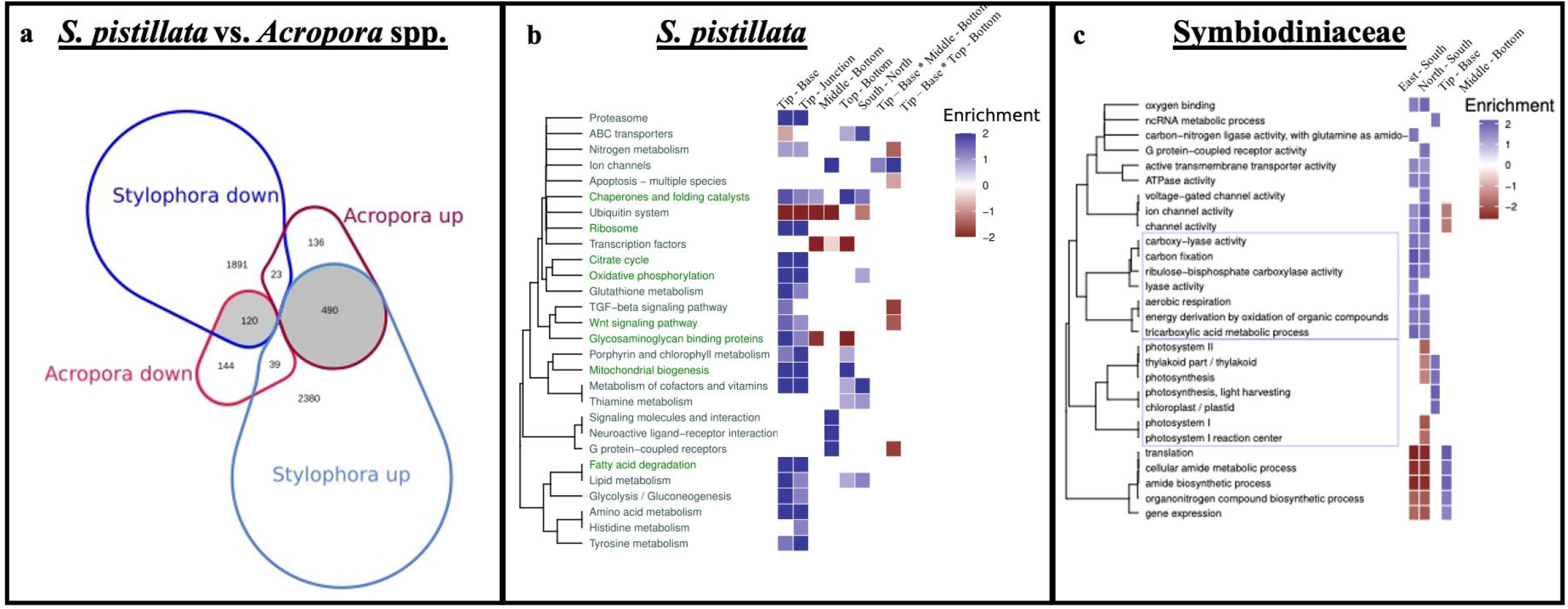
Differential gene expression across coral species and terms clustering in host and symbiont. Venn diagram (a) representing the proportional counts of shared tip vs. base DE genes in *S. pistillata* (here) and two species of *Acropora* (Hemond et al., 2014) (Supplementary Table 9), with shaded regions representing genes showing the same DE patterns and noting “up” and “down” regulated genes. Enrichment patterns of clustered host KEGG (*S. pistillata;* b) and photosymbiont GO terms (UniProt *S. microadriaticum*; c) across factor comparisons. Host terms enriched in both *S. pistillata* (here) and two species of *Acropora* (Hemond et al., 2014) for tip vs base comparisons, are shown as green text. Hierarchical tree clustering reflects functional similarity between terms. Color-coded heatmap scores equal to log_10_(adjusted p-value) of enriched terms multiplied by 1 and −1 for up- and down-regulated genes, respectively (for clarity, values <-2 and >2 are shown as 2 and −2).

In contrast to the host, of ~12,000 tested photosymbiont genes, only ~150 were differentially expressed (DE) (Supplementary Table 7). The host vs. symbiont differences are partly attributed to technical differences in statistical power due to lower symbiont read counts, and likely represent an underestimation of symbiont DE genes. DE gene patterns of the photosymbionts are not as clearly delineated as for the host (Figure 2d); nevertheless, the expression patterns differ between cardinal directions (p<0.0007, ADONIS test, Supplementary Table 8), likely reflecting the different light regimes experienced on different sides of the colony, visible as shading on the north (right) side of the colony and no shading on the south (left) side of the colony in Figure 1a, although we did not measure light levels around the colony. The ultimate drivers of the differences observed for groups of polyps in locations differing by ring height and cardinal direction remain to be resolved. Further, the observed differences across a single colony could have implications for studies using branches collected from different locations on separate colonies. While intracolonial differences that may diminish observable transcriptional and physiological differences between treatments - if chosen branches are from differing parts of selected colonies - may lessen over multi-year studies (e.g., Palumbi et al., 2014), epigenetic responses that differ across a colony may persist across generations (Liew et al., 2020).

### 4.3 Patterns in gene expression are conserved between coral taxa

A previous study, analyzing the differences in gene expression between branch tips and bases in *Acropora* spp. (Hemond et al., 2014), provided an opportunity to identify conserved intra-colonial patterns in gene expression between these two coral taxa, which likely diverged at least 200 million years ago (Park et al., 2012). We identified ~9,600 putative orthologous expressed genes between *S. pistillata* and two species of *Acropora* spp. (Hemond et al., 2014), based on reciprocal BLAST. Interestingly, from a total of 952 genes that were DE in *Acropora* spp. along the branch axis, ~70% were also DE in *S. pistillata* (672), and ~90% of these exhibited the same fold change trend (p-value < 2.2e-16, Fisher’s exact test; Figure 3a, Supplementary Table 9). Overall, these results indicate that micro-environmental dependence of gene differential expression at tips versus bases reflects an evolutionarily conserved pattern rather than a species-specific pattern, being observed in members of both robust (here) and complex (Hemond et al., 2014) scleractinian clades, as well as between taxa which possess physiologically unique axial polyps (*Acropora;* (Wallace, 1978;1985) and a genus which does not (*Stylophora)*. It also supports that the occurrence of intracolonial differential responses to microenvironments in general, and up-regulation of genes for specific processes such as protein production and energy generation in particular (Figure 3b, green font), can likely be generalized across branching coral taxa.

### 4.4 Gene expression patterns suggest differences in host versus photosymbiont metabolism

We next inferred potential broad-scale changes in cellular and metabolic functions by identifying KEGG and Gene Ontology (GO) enrichment patterns associated with micro-environmental changes in both the host (Figure 3b, Supplementary Tables 10, 11) and the photosymbiont (Figure 3c, Supplementary Table 12). As expected, broad clusters of host KEGG terms and photosymbiont GO terms were DE (Figure 3b, c), mirroring the pattern observed for all DE genes within the host and symbiont (Figure 2c, d), with many more terms DE in the coral host along the branch axis compared to the ring height or cardinal direction (Figure 3b), and more terms differed along cardinal direction for the photosymbiont (Figure 3c).

Most DE host genes grouped by KEGG term tended to be up-regulated in branch tips relative to the junctions or bases (Figure 3b, Supplementary Tables 10, 11). Specifically, many terms related to cell metabolism were enriched in the tips versus the bases, including carbohydrate metabolism (e.g., glycolysis and the TCA cycle, oxidative phosphorylation, mitochondrial biosynthesis), amino acid biosynthesis and metabolism, and lipid metabolism (Figure 3b). We further tested which functional groups were enriched with conserved tip vs. base DE gene patterns between coral taxa (Figure 3a, b). As indicated by green KEGG term text in Figure 3b, many fundamental processes such as metabolism show similar DE trends and enrichment between these two genera (Supplementary Tables 9, 10), indicating a conserved response between the taxa. “Zooming in” from the KEGG terms to the underlying genes, we note that genes related to oxidative phosphorylation, such as ATP synthase, cytochrome c oxidase, co-enzyme Q, and NADH dehydrogenase tend to be up-regulated at branch tips across the species. In addition, multiple genes for succinate dehydrogenase as well as other individual genes comprising the citric acid cycle, were also more highly expressed at branch tips than junctions or bases in *S. pistillata* and many of these showed similar regulation patterns in the *Acropora* spp. Finally, most of the >100 DE genes with KEGG terms for mitochondrial biogenesis were up-regulated at tips compared with junctions and bases in *S. pistillata* and many of their likely homologs were also up-regulated at branch tips in *Acropora* spp. The increased rates in physiological processes involved in energy production at branch tips, as suggested by our gene expression analysis, have been observed in the branching coral *A. palmata* as increased respiration and mitotic index (Gladfelter et al., 1989).

Intriguingly, the patterns of gene expression observed in the host were not all recapitulated in the photosymbiont. Although some symbiont GO terms associated with photosynthesis were enriched in the tips vs bases in the symbionts (e.g., those involved in light-harvesting and the thylakoid), likely in response to higher light intensities at tips compared to bases (e.g., Helmuth et al., 1997;Kaniewska et al., 2008), others were not, including terms involved with carbon fixation (Figure 3c). Additionally, no differing patterns in photophysiology were found along branch axis, measured here as chlorophyll and symbiont density per both total protein and area (Supplementary Tables 2, 13, Supplementary Figure 6), although we acknowledge that these are limited measures of the photophysiological processes. Indeed, previous studies have shown that the products of photosynthesis can be translocated from branch bases to the tips in branching corals (e.g., Rinkevich and Loya). Rather than exhibiting tip-to-base functional differences, many of the terms associated with symbiont photosynthesis (both light and dark reactions) were found to differ along cardinal direction, being enriched in south-facing branches (Figure 3c), possibly due to the direction of the sun. As such, we anticipate that future studies will find differences in gene expression in the symbiont across a diel cycle, with, for example potentially decreased differences between north- and south-facing branches during nighttime hours. As noted above, these enrichment patterns are likely an underestimation. Furthermore, correlations between mRNA abundance and physiological patterns may not be high due to translation level effects, translocation of host/symbiont products, food particle capture by the host, etc., which may lead to differences in symbiont transcriptional control.

### 4.5 Developmental pathways are DE along the branch axis

In addition to pathways related to metabolism, several KEGG terms associated with development and morphological plasticity in colony architecture were DE between branch tips and junctions or bases. For example, the Wnt signaling pathway was enriched at branch tips versus the bases in both *S. pistillata* (Figure 3b) and *Acropora* spp. (Hemond et al., 2014). This pathway, which transmits information from outside to inside cells, is associated with cell differentiation and axis patterning in cnidarians (reviewed in Lee et al., 2006;e.g., Duffy et al., 2010) and bilaterians (e.g., Siegfried and Perrimon). Similarly, the TGFβ signaling pathway was also enriched in *S. pistillata* branch tips (Figure 3b), although this was not observed in *Acropora* spp. (Hemond et al., 2014); this pathway has been implicated for a role in development and calcification in corals (Gutner-Hoch et al., 2017). At the individual gene level, we observed that fibroblast growth factors, which could aid in development of limbs (reviewed in Xu et al., 1999) and in production of nerve cells (reviewed in Guillemot and Zimmer, 2011), were significantly up-regulated at branch tips. Half of the genes involved in mitochondrial biosynthesis were also up-regulated at branch tips. Further, two different bone morphogenetic proteins (BMPs), which may play a role in axial patterning in cnidarian polyps (Reinhardt et al., 2004) and in budding (Surekha et al., 2020), were up-regulated at branch tips here. Finally, several retinol dehydrogenases, proteins associated with photoreceptors in higher organisms (Albalat, 2012), were up-regulated at branch tips, where light levels tend to be higher, an observation shared with *Acropora* spp. (Hemond et al., 2014).

In addition to the plasticity described above, gamete production has also been shown to vary along branches axes across several coral taxa with gametogenesis typically occurring preferentially deeper in the colony (e.g., Stimson). We observed the highest expression of several testis-specific genes, such as testis-specific chromodomain protein Y 1-like and testis-specific protein kinases at *S. pistillata* branch junctions (Supplementary Table 4). In humans, these genes are associated with chromosome Y (Lahn and Page) and/or may be involved in sperm structure (Walden and Cowan). Interestingly, we observed up-regulation of octopamine receptors at branch tips, suggestive of higher maturation of ovaries and testes in that location (Chiu et al., 2020).

### 4.6 Changes in expression of genes potentially related to predation along branch axis

As described above, many metabolic processes were enriched in the tips versus the bases, whereas there is little evidence for higher photosynthesis by photosymbionts at branch tips. Asking whether this could be due to more heterotrophic food input in the tips (e.g., through predation), we analyzed the tissue toxicity (hemolytic activity) and gene expression patterns of putative toxins. Although there was no difference in nematocyst abundance along branch axes (Figure 4a, Supplementary Table 2), tissue hemolytic activity was highest at branch tips (Figure 4b, Supplementary Table 2), consistent with previous studies in *S. pistillata* that used branches from two different colonies (Ben-Ari et al.). Thus, the observed branch axis patterning of hemolysis potential is robust across at least three colonies from two independent studies. Using reciprocal BLAST, we identified 22 putative toxin genes (Rachamim et al., 2015), encoding cytolysins, phospholipase A2 toxins, SCRiPs (small cysteine rich peptides) and a potential Kunitz/K channel toxin (Supplementary Table 14). We also compared these genes to the reciprocal BLAST results against the *Acropora* spp. dataset (Hemond et al., 2014), finding only two orthologous matches, a SCRiP and an uncharacterized protein. Like their *S. pistillata* counterparts, they were not DE between branch tips and bases (Supplementary Table 15). In our putative toxin dataset, we observed 13 and 11 DE genes between branch tips versus bases or junctions, respectively (Figure 4c, d; Supplementary Table 15, Supplementary Figure 7). Up-regulation of genes encoding potential toxins at branch tips has previously been observed in colonial corals (Hemond et al.;Hemond and Vollmer). The two most abundantly expressed putative toxin genes encode for a cytolysin (up-regulated at tips) and a second SCRiP (up-regulated at bases), with the rest of the toxin genes having much lower overall expression levels (Figure 4c, d; Supplementary Table 15). The highly expressed cytolysin is of particular interest as it belongs to the actinoporin family, and a hemolytic protein from this family, likely the protein or an isoform of the highly expressed gene here (Supplementary Figure 8), has previously been isolated from *S. pistillata* (Ben-Ari et al., 2018). Actinoporins are ~20 kDa water-soluble proteins which are lethal when injected into crustaceans (Giese et al., 1996), and which insert themselves into membranes, particularly those containing sphingomyelins (Alvarez et al., 2009). They have been immunolocalized to tentacle nematocysts as well as to mesenterial filaments (Basulto et al., 2006). SCRiPs are ~5 kDa peptides which may be anthozoan-specific (Sunagawa et al.;Jouiaei et al., 2015), with evidence that they have neurotoxic effects (Jouiaei et al., 2015), although their function remains to be fully clarified.

**Figure 4.**
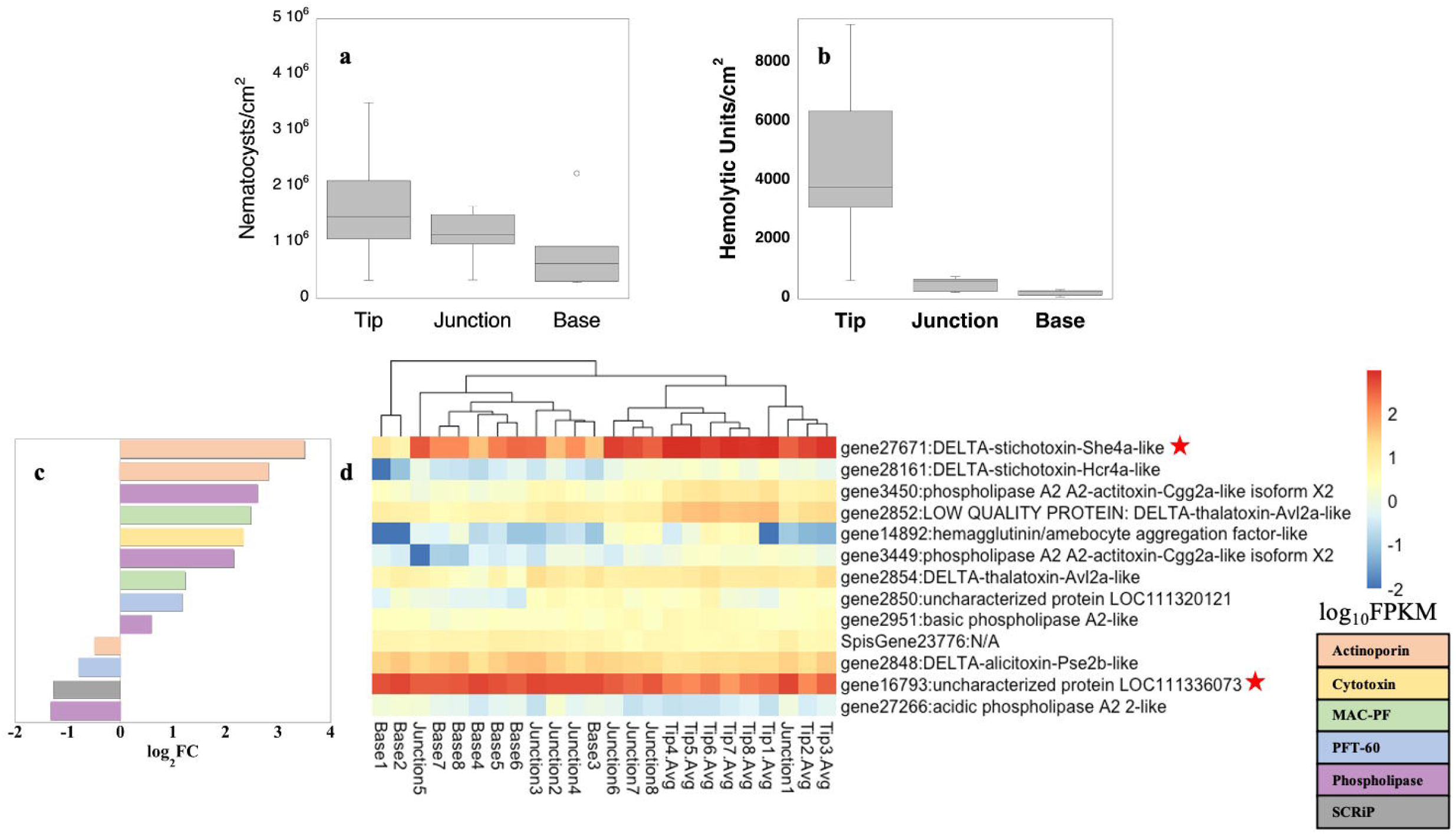
Toxicity potential. Nematocyst abundance (a) and hemolytic units per area (b), and gene expression patterns of putative toxins as log2FC between branch tips (positive values) and bases (negative values) (c), and log_10_FPKM along *S. pistillata* branch axis (d). Only significantly DE toxin genes are shown (p<0.05). The two most abundant transcripts, a cytolysin annotated as stichotoxin and an uncharacterized gene, for which reciprocal BLAST hits suggest it is a SCRiP (Supplementary14 Table), are noted with a red star (d).

The higher hemolytic activity and larger number of toxins more abundantly expressed at branch tips suggest that the tips play the dominant role in venom-based competition, defense, and predation, although more work is needed to identify the role of putative toxins such as SCRiPs, which were more abundantly expressed at branch bases, and the potential contribution and roles of photosymbiont toxin production (e.g., Nakamura et al., 1993). While it is difficult to disentangle toxin effects on competition and defense from predation, based on higher flow at the edges of colonies (Chang et al., 2009), colonies’ edges are likely exposed to higher food loads than are areas closer to the base of the branches. Consumption of these food sources could then be reflected in increased respiration rates as the food is metabolized, as has been observed in other species (Gladfelter et al., 1989) and may yield increased energy production which would require up-regulation of genes involved in this process as we observed here. We also queried genes related to digestion and observed that many peptidases were DE, but not in a clear pattern along the branches with approximately half up-regulated at the tips. Similarly, multiple lipases were DE between branch locations, but no general trend was observed. However, pancreatic lipase expression, which has been observed in gastric cirri in a scyphozoan (Steinmetz, 2019), was found to be significantly up-regulated to a high degree at branch tips. We also observed up-regulation of three chitinases in branch tips versus bases, which could support extracellular digestion of arthropod prey (Steinmetz, 2019). In contrast, an amylase which converts starch and glycogen - biomolecules used by the photosymbiont and host, respectively, to store photosynthates (Kopp et al., 2015) - into simple sugars, was down-regulated at the tips versus junctions and bases. Thus, while our results suggest differences along the branch axis of functions related to prey capture and defense, including both toxicity and digestion, further work is needed to fully characterize the relative contributions of photosynthesized products and predation. Additionally, temporal differences in predation capabilities at both the transcriptional (i.e., genes related to digestion) and physiological (i.e., hemolytic potential) levels across individual colonies should be examined as zooplankton abundance on coral reefs changes on a diel cycle (Yahel et al., 2005). Future studies measuring water flow rates explicitly in conjunction with transcriptional changes across colonies with complex macromorphologies will also clarify precisely how particle capture and digestion are affected by differing flow patterns (e.g., Sebens et al., 1997).

### 4.7 Changes in skeleton structure and expression of known biomineralization genes along branch axis

Shallow water stony corals are most notable for their production of massive reef structures, growing in size by adding mineral and new polyps, particularly for branching corals, at colony tips or edges (e.g., D’A and Le Tissier, 1988;Meesters and Bak, 1995). We therefore examined in depth skeletal morphological patterns and related gene expression across the three branch axis locations. Unlike most other physiological parameters across the colony which were not significantly different between branch positions (Supplementary Tables 2, 13), those linked to differences in biomineralization were significantly different along branch axes. Qualitatively, coenosteal spines appear narrowest and thinnest at the tip while base spines exhibit a more distinct microtuberculate texture in their lower portions (Figure 5aii, iv, and vi). Such microtuberculate texture with better differentiation of mineral fiber bundles may suggest higher organic matrix content which separates the mineral phase packages to a greater extent (Mass et al., 2014;Coronado et al., 2019). Corallites were closer together and more elliptically shaped at the tips than at the branch junctions or bases (Figure 5b and c; Supplementary Tables 2, 13). Comparable trending toward ellipticity at branch tips has been reported in other Pocilloporids (Schmidt-Roach et al., 2014). Further, genetically identical *S. pistillata* branches have been shown to have more distantly spaced corallites with thicker theca and longer septa when grown under light spectra biased toward blue wavelengths (Rocha et al., 2014). While we did not measure light spectra in this study, such a spectral shift along a single branch, due to absorbance of red light by water, has been observed in *S. pistillata* tissue at the colony level on the sides versus at the top of ~30 cm diameter colonies (Kaniewska et al., 2011) and at the branch level along *Acropora millepora* branches toward the colony interior (Wangpraseurt et al., 2014).

**Figure 5.**
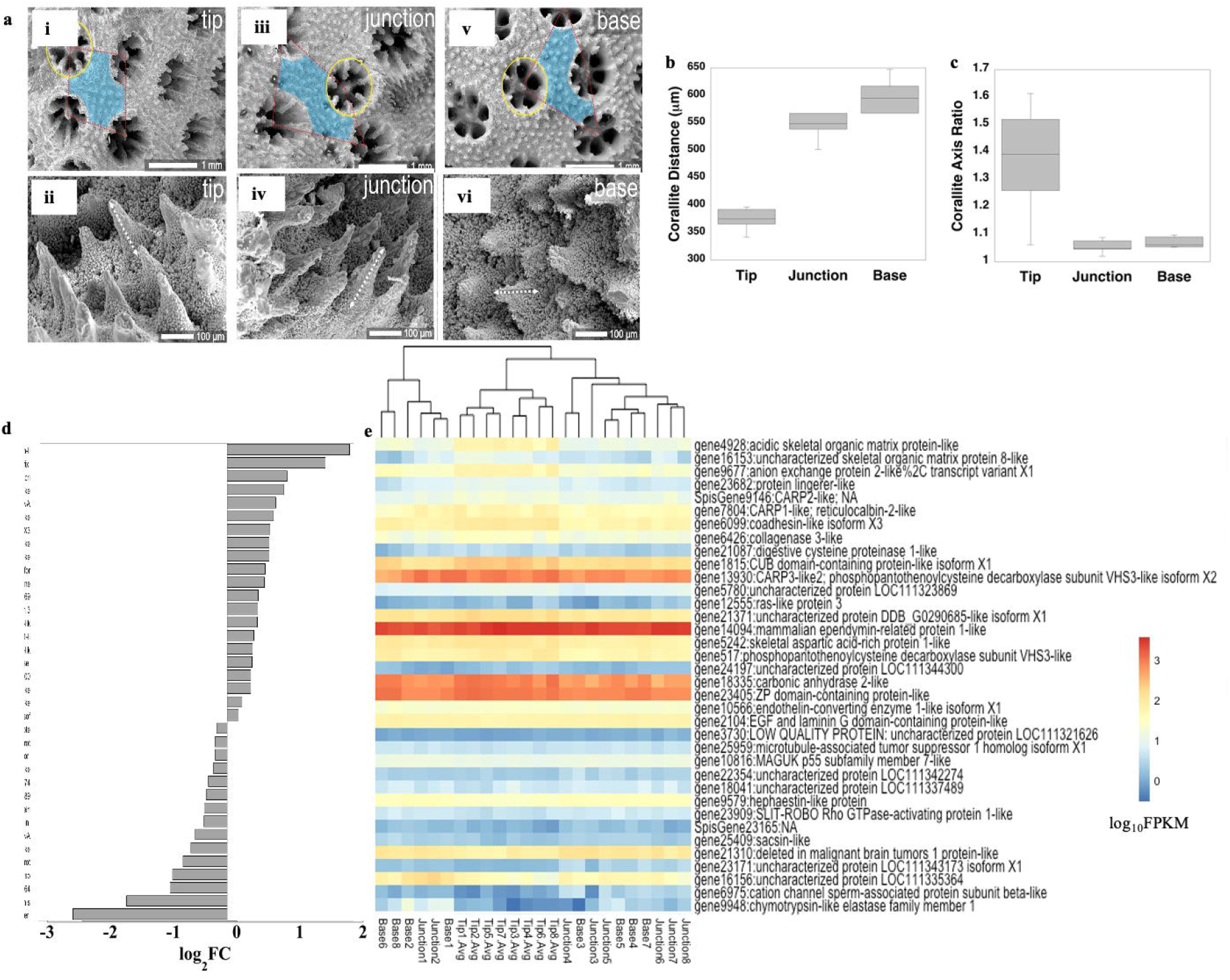
Skeleton characteristics and biomineralization gene expression. Microstructural differences of corallites and coenosteal spines along the branch axis (a; tip in i, ii; junction in iii, iv; base in v, vi). Corallite spacing (b) and ellipticity (c) along *S. pistillata* branch axis. Several known biomineralization genes were DE, shown as log2FC between branch tips (positive values) and bases (negative values) (d) and log_10_FPKM along the branch axis (e) to clarify genes with high versus low expression. Only significantly DE genes are shown (p<0.05).

We next examined differential expression of genes encoding proteins sequenced from *S. pistillata* skeleton and likely directly involved in the calcification mechanism (Puverel et al., 2005;Bertucci et al., 2011;Drake et al., 2013;Mass et al., 2014;Peled et al., 2020;Mummadisetti et al., 2021), as well as a bicarbonate anion transporter observed predominantly in the calicoblastic cell layer (Zoccola et al., 2015) (Figure 5e; Supplementary Table 16, Supplementary Figure 9). We did not include genes for known skeletal proteins from *Acropora millepora* and *A. digitifera* (Ramos-Silva et al.;Takeuchi et al.), as not all orthologs have been found in *S. pistillata* skeleton although they exist in the genome, or non-orthologous proteins within the same functional group occur in *S. pistillata* skeleton (Zaquin et al., 2021). Similar to the global gene expression pattern for the host, we observed an up-regulation of biomineralization genes biased toward the tips (Figure 5d, e; Supplementary Table 16). Of the 36 biomineralization-related genes that were DE between branch tips and bases, eight orthologous genes were also DE and in the same direction in *Acropora* spp. (Hemond et al., 2014) (Supplementary Table 17). In *S. pistillata*, the bicarbonate anion transporter SLCg4, a coadhesin-like protein, uncharacterized skeletal organic matrix protein 8 (USOMP8), and a hypothetical protein were up-regulated at branch tips but were expressed in only low amounts. In contrast, a ZP-domain containing protein, carbonic anhydrase (STPCA2), mammalian ependymin-related protein, and coral acid rich protein 3 (CARP3) were moderately up-regulated at branch tips and were very highly expressed overall. These proteins exhibit very different functions with SLCg4 and STPCA2 involved in regulating carbonate chemistry within the calcifying space (Zoccola et al., 2015;Zoccola et al., 2016), while CARP3, which is a highly acidic protein (35% Asp, 15%Glu (Mass et al., 2013)) can direct mineralogy toward formation of Mg-calcite and vaterite (Gavriel et al., 2018;Laipnik et al., 2020), and is up-regulated in newly-settled coral polyps (Akiva et al., 2018), in colonies growing at shallow depths (Malik et al., 2021), and in coral cell cultures reared at increased pCO2 (Drake et al., 2017). Although the function of USOMP8 is not known, it is structurally similar to a BMP inhibitor (Drake et al., 2013). Of the remaining known highly acidic coral skeleton genes, CARPs 1 and 4 (CARP4 is sometimes also called skeletal aspartic acid rich protein 1; SAARP1 (Ramos-Silva et al., 2013)) were up-regulated at branch tips and were moderately expressed; both proteins are biased toward Asp residues (Mass et al., 2013) and are likely involved in nucleation initiation in early mineralization zones of growing septa (Mass et al., 2014). In contrast, CARP2 exhibited very low although significantly up-regulated expression at branch tips; while this protein is suggested to have a role in skeleton extension based on its localization to more soluble growth areas (Mass et al., 2014), its down-regulation upon settlement in pocilloporid larvae (Mass et al., 2016;Akiva et al., 2018), association with ACC phases pre-settlement (Akiva et al., 2018), and low overall expression in the present study may instead point to a role in inhibiting aragonite nucleation.

Previous and our present work provide strong genetic support for highest biomineralization activity at branch tips (e.g., Goreau, 1959). This faster calcification at the tips may result in the qualitatively thinner and narrower coenosteal spines we observed at branch tips (Figure 5) that would be thickened by polyps as they age. Energy for higher calcification rates at branch tips could come from two sources. First, there is evidence of translocation of ATP from tissue regions deeper in the colony with relatively high photosynthetic rates (Fang et al., 1989) to branch tips. Secondly, our and previous data suggest that the potential for predation is higher at branch tips, with higher toxin gene expression and hemolytic activity at that location (Figure 4) (Hemond et al., 2014;Ben-Ari et al., 2018), and up-regulation of genes related to metabolism and energy production (Figure 3). This further suggests that biomineralization gene expression and skeleton morphology patterns observed at the sub-polyp scale (i.e., centers of calcification in growing tips of septa) are recapitulated at the branch scale (Malik et al., 2021).

## 5 Summary and Conclusions

Some colonial cnidarians, particularly hydrozoans, exhibit a clear division of labor inside the colony, with functionally and morphologically distinct polyps across the colony (Cartwright et al., 1999). While no obvious division of labor at the polyp scale is observed in corals, our results show, from molecular to physiological and morphological scales on a single genetically-identical colony, a functional specialization by clusters of polyps due to their locations on coral branches and around the colony likely related to the microenvironments they encounter (Rueffler et al., 2012) (Figure 6). In simplest terms, polyps at branch tips and toward the top of the colony are more exposed to light and higher water flow than are those found lower in the branches and closer to the substrate with consequent differential expression of a multitude of genes directing many key functions and, at least along the branch axis, differences in physiology and skeleton morphology. Our findings suggest that branch tips may also be sites of higher potential for envenomation for colony defense and predation, as well as possibly for enhanced prey capture. These plastic responses by genetically identical polyps can then be communicated through the coenosarc to the rest of the colony to allow for enhanced resource sharing (Mackie, 1986). Such plasticity may aid in colony recovery after partial mortality events, such as those due to bleaching (Matsuda et al., 2020), and may be integral to adaptation to environmental change (Kenkel and Matz, 2016). Further, the expression patterns underlying the tested micro-environmental differences are found to be evolutionary conserved in the distantly-related stony corals *S. pistillata* and *Acropora* spp. Future study of multiple colonies and species will clarify the extent to which the general patterns we observe across the colony studied here are found across differing colony morphologies, diurnal and seasonal cycles, and geographic location. Further, the diversity of gene expression and physiology across a coral colony needs to be taken into account when designing experiments with coral fragments from multiple colonies.

**Figure 6.**
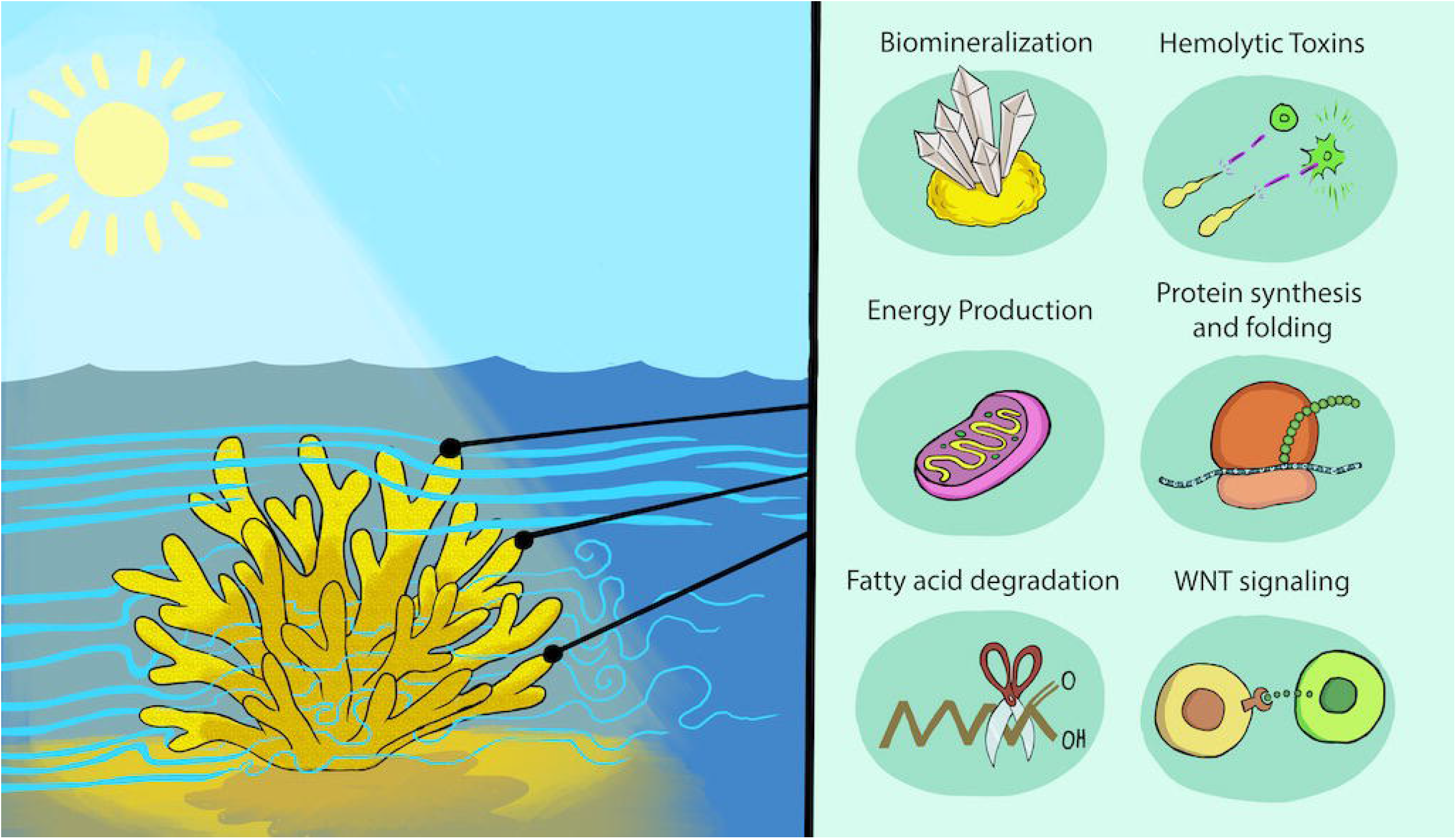
Micro-environmental variation across a colony leads to physiological and transcriptional differences. Schematic of environmental parameters (e.g., flow and light intensity, left) and resulting differential gene expression patterns and physiological outcomes for key processes focusing on those enhanced at branch tips (right). We cannot assign gene expression to specific tissue layers or cell types in most cases, except for the likely localization of toxin and biomineralization genes to the oral ectoderm and calicoderm, respectively.

## 8 Author Contribution Statement

YP, OY, DT, DS, and TM conceived the study. YP, OY, ES prepared the samples. JLD, AM, YP, OY, ES, and JS conducted formal analysis of the data. All authors wrote the manuscript and approve of this submission.

## 10 Acknowledgements

We thank Yarden Ben-Tabou de-Leon for drawing the summary Figure 6 and Dikla Aharonovich for guidance and laboratory assistance. We thank the Crown Genomics Institute of the Nancy and Stephen Grand Israel National Center for Personalized Medicine, Weizmann Institute of Science for transcriptome sequencing.

## 11 Data Availability Statement

The data underlying this article are available in the article, in its online Supplementary Material, and on GitHub in the following repository: https://github.com/jeanadrake/Single_coral_colony_transcriptomes_physiology.

## References

Akiva, A., Neder, M., Kahil, K., Gavriel, R., Pinkas, I., Goobes, G., and Mass, T. (2018). Minerals in the pre-settled coral *Stylophora pistillata* crystallize via protein and ion changes. Nature Communications 9, 1880.

Albalat, R. (2012). Evolution of the genetic machinery of the visual cycle: a novelty of the vertebrate eye? Molecular Biology and Evolution 29, 1461–1469.

Alvarez, C., Mancheno, J.M., Martínez, D., Tejuca, M., Pazos, F., and Lanio, M.E. (2009). Sticholysins, two pore-forming toxins produced by the Caribbean sea anemone *Stichodactyla helianthus:* their interaction with membranes. Toxicon 54, 1135–1147.

Anthony, K.R., Hoogenboom, M.O., and Connolly, S.R. (2005). Adaptive variation in coral geometry and the optimization of internal colony light climates. Functional Ecology 19, 17–26.

Aranda, M., Li, Y., Liew, Y.J., Baumgarten, S., Simakov, O., Wilson, M.C., Piel, J., Ashoor, H., Bougouffa, S., and Bajic, V.B. (2016). Genomes of coral dinoflagellate symbionts highlight evolutionary adaptations conducive to a symbiotic lifestyle. Scientific Reports 6, 1–15.

Bartosz, G., Finkelshtein, A., Przygodzki, T., Bsor, T., Nesher, N., Sher, D., and Zlotkin, E. (2008). A pharmacological solution for a conspecific conflict: ROS-mediated territorial aggression in sea anemones. Toxicon 51, 1038–1050.

Basulto, A., Pérez, V.M., Noa, Y., Varela, C., Otero, A.J., and Pico, M.C. (2006). Immunohistochemical targeting of sea anemone cytolysins on tentacles, mesenteric filaments and isolated nematocysts of *Stichodactyla helianthus*. Journal of Experimental Zoology Part A: Comparative Experimental Biology 305, 253–258.

Ben-Ari, H., Paz, M., and Sher, D. (2018). The chemical armament of reef-building corals: inter- and intra-specific variation and the identification of an unusual actinoporin in *Stylophora pistilata*. Scientific Reports 8, 1–13.

Bertucci, A., Tambutté, S., Supuran, C., Allemand, D., and Zoccola, D. (2011). A new coral carbonic anhydrase in *Stylophora pistillata*. Marine Biotechnology 13, 992–1002.

Bryant, D.M., Johnson, K., DiTommaso, T., Tickle, T., Couger, M.B., Payzin-Dogru, D., Lee, T.J., Leigh, N.D., Kuo, T.-H., and Davis, F.G. (2017). A tissue-mapped axolotl de novo transcriptome enables identification of limb regeneration factors. Cell Reports 18, 762–776.

Buchfink, B., Xie, C., and Huson, D.H. (2015). Fast and sensitive protein alignment using DIAMOND. Nature Methods 12, 59–60.

Carpenter, L.W., and Patterson, M.R. (2007). Water flow influences the distribution of photosynthetic efficiency within colonies of the scleractinian coral *Montastrea annularis* (Ellis and Solander, 1786); implications for coral bleaching. Journal of Experimental Marine Biology and Ecology 351, 10–26.

Cartwright, P., Bowsher, J., and Buss, L.W. (1999). Expression of a Hox gene, Cnox-2, and the division of labor in a colonial hydroid. Proceedings of the National Academy of Sciences 96, 2183–2186.

Chang, S., Elkins, C., Alley, M., Eaton, J., and Monismitha, S. (2009). Flow inside a coral colony measured using magnetic resonance velocimetry. Limnology and Oceanography 54, 1819–1827.

Chiu, Y.-L., Shikina, S., Yoshioka, Y., Shinzato, C., and Chang, C.-F. (2020). De novo transcriptome assembly from the gonads of a scleractinian coral, *Euphyllia ancora:*molecular mechanisms underlying scleractinian gametogenesis. BMC Genomics 21, 1–20.

Conlan, J.A., Humphrey, C.A., Severati, A., and Francis, D.S. (2018). Intra-colonial diversity in the scleractinian coral, *Acropora millepora*: identifying the nutritional gradients underlying physiological integration and compartmentalised functioning. PeerJ 6, e4239.

Coronado, I., Fine, M., Bosellini, F.R., and Stolarski, J. (2019). Impact of ocean acidification on crystallographic vital effect of the coral skeleton. Nature Communications 10, 1–9.

D’A, M., and Le Tissier, A. (1988). The growth and formation of branch tips of *Pocillopora damicornis* (Linnaeus). Journal of Experimental Marine Biology and Ecology 124, 115–131.

Dennison, W.C., and Barnes, D.J. (1988). Effect of water motion on coral photosynthesis and calcification. Journal of Experimental Marine Biology and Ecology 115, 67–77.

DePristo, M.A., Banks, E., Poplin, R., Garimella, K.V., Maguire, J.R., Hartl, C., Philippakis, A.A., Del Angel, G., Rivas, M.A., and Hanna, M. (2011). A framework for variation discovery and genotyping using next-generation DNA sequencing data. Nature Genetics 43, 491.

Dobin, A., Davis, C.A., Schlesinger, F., Drenkow, J., Zaleski, C., Jha, S., Batut, P., Chaisson, M., and Gingeras, T.R. (2013). STAR: ultrafast universal RNA-Seq aligner. Bioinformatics 29, 15–21.

Drake, J.L., Mass, T., Haramaty, L., Zelzion, E., Bhattacharya, D., and Falkowski, P.G. (2013). Proteomic analysis of skeletal organic matrix from the stony coral *Stylophora pistillata*. Proceedings of the National Academy of Sciences 110, 3788–3793.

Drake, J.L., Schaller, M.F., Mass, T., Godfrey, L., Fu, A., Sherrell, R.M., Rosenthal, Y., and Falkowski, P.G. (2017). Molecular and geochemical perspectives on the influence of CO2 on calcification in coral cell cultures. Limnology and Oceanography.

Duffy, D.J., Plickert, G., Kuenzel, T., Tilmann, W., and Frank, U. (2010). Wnt signaling promotes oral but suppresses aboral structures in *Hydractinia* metamorphosis and regeneration. Development 137, 3057–3066.

Falkowski, P.G., Dubinsky, Z., Muscatine, L., and Porter, J.W. (1984). Light and the bioenergetics of a symbiotic coral. Bioscience 34, 705–709.

Fang, L.-S., Chen, Y.-w.J., and Chen, C.-S. (1989). Why does the white tip of stony coral grow so fast without zooxanthellae? Marine Biology 103, 359–363.

Garren, M., Son, K., Raina, J.-B., Rusconi, R., Menolascina, F., Shapiro, O.H., Tout, J., Bourne, D.G., Seymour, J.R., and Stocker, R. (2014). A bacterial pathogen uses dimethylsulfoniopropionate as a cue to target heat-stressed corals. The ISME Journal 8, 999–1007.

Gavriel, R., Nadav-Tsubery, M., Glick, Y., Yarmolenko, A., Kofman, R., Keinan-Adamsky, K., Berman, A., Mass, T., and Goobes, G. (2018). The coral protein CARP3 acts from a disordered mineral surface film to divert aragonite crystallization in favor of Mg-calcite. Advanced Functional Materials 28, 1707321.

Gibbin, E., Gavish, A., Domart-Coulon, I., Kramarsky-Winter, E., Shapiro, O., Meibom, A., and Vardi, A. (2018). Using NanoSIMS coupled with microfluidics to visualize the early stages of coral infection by *Vibrio coralliilyticus*. BMC microbiology 18, 1–10.

Giese, C., Mebs, D., and Werding, B. (1996). Resistance and vulnerability of crustaceans to cytolytic sea anemone toxins. Toxicon 34, 955–958.

Gladfelter, E.H., Michel, G., and Sanfelici, A. (1989). Metabolic gradients along a branch of the reef coral *Acropora palmata*. Bulletin of Marine Science 44, 1166–1173.

Goreau, T.F. (1959). The physiology of skeleton formation in corals. I. A method for measuring the rate of calcium deposition by corals under different conditions. Biological Bulletin 116, 59–75.

Guillemot, F., and Zimmer, C. (2011). From cradle to grave: the multiple roles of fibroblast growth factors in neural development. Neuron 71, 574–588.

Gutner-Hoch, E., Ben-Asher, H.W., Yam, R., Shemesh, A., and Levy, O. (2017). Identifying genes and regulatory pathways associated with the scleractinian coral calcification process. PeerJ 5, e3590.

Helmuth, B.S., Timmerman, B., and Sebens, K. (1997). Interplay of host morphology and symbiont microhabitat in coral aggregations. Marine Biology 130, 1–10.

Hemond, E.M., Kaluziak, S.T., and Vollmer, S.V. (2014). The genetics of colony form and function in Caribbean *Acropora* corals. BMC Genomics 15, 1133.

Hemond, E.M., and Vollmer, S.V. (2015). Diurnal and nocturnal transcriptomic variation in the Caribbean staghorn coral, *Acropora cervicornis*. Molecular Ecology 24, 4460–4473.

Hoegh-Guldberg, O., Jacob, D., Taylor, M., Bolaños, T.G., Bindi, M., Brown, S., Camilloni, I., Diedhiou, A., Djalante, R., and Ebi, K. (2019). The human imperative of stabilizing global climate change at 1.5° C. Science 365, eaaw6974.

Hossain, M.M., and Staples, A.E. (2020). Mass transport and turbulent statistics within two branching coral colonies. Fluids 5, 153.

Hughes, T.P. (1994). Catastrophes, phase shifts, and large-scale degradation of a Caribbean coral reef. Science 265, 1547–1551.

Jouiaei, M., Sunagar, K., Federman Gross, A., Scheib, H., Alewood, P.F., Moran, Y., and Fry, B.G. (2015). Evolution of an ancient venom: recognition of a novel family of cnidarian toxins and the common evolutionary origin of sodium and potassium neurotoxins in sea anemone. Molecular Biology and Evolution 32, 1598–1610.

Kaniewska, P., Anthony, K.R., and Hoegh-Guldberg, O. (2008). Variation in colony geometry modulates internal light levels in branching corals, *Acropora humilis* and *Stylophora pistillata*. Marine Biology 155, 649–660.

Kaniewska, P., Magnusson, S.H., Anthony, K.R., Reef, R., Kühl, M., and Hoegh-Guldberg, O. (2011). Importance of macro-versus microstructure in modulating light levels inside coral colonies. Journal of Phycology 47, 846–860.

Kemp, D.W., Hernandez-Pech, X., Iglesias-Prieto, R., Fitt, W.K., and Schmidt, G.W. (2014). Community dynamics and physiology of Symbiodinium spp. before, during, and after a coral bleaching event. Limnology and Oceanography 59, 788–797.

Kenkel, C.D., and Matz, M.V. (2016). Gene expression plasticity as a mechanism of coral adaptation to a variable environment. Nature Ecology & Evolution 1, 1–6.

Knowlton, N., Brainard, R.E., Fisher, R., Moews, M., Plaisance, L., and Caley, M.J. (2010). Coral reef biodiversity. Life in the world’s oceans: diversity distribution and abundance, 65–74.

Kopp, C., Domart-Coulon, I., Escrig, S., Humbel, B.M., Hignette, M., and Meibom, A. (2015). Subcellular investigation of photosynthesis-driven carbon assimilation in the symbiotic reef coral *Pocillopora damicornis*. MBio 6.

Lahn, B.T., and Page, D.C. (1999). Retroposition of autosomal mRNA yielded testis-specific gene family on human Y chromosome. Nature Genetics 21, 429–433.

Laipnik, R.a., Bissi, V., Sun, C.-Y., Falini, G., Gilbert, P.U., and Mass, T. (2020). Coral acid rich protein selects vaterite polymorph in vitro. Journal of Structural Biology 209, 107431.

LaJeunesse, T.C., Parkinson, J.E., Gabrielson, P.W., Jeong, H.J., Reimer, J.D., Voolstra, C.R., and Santos, S.R. (2018). Systematic revision of Symbiodiniaceae highlights the antiquity and diversity of coral endosymbionts. Current Biology 28, 2570–2580. e2576.

Lee, P.N., Pang, K., Matus, D.Q., and Martindale, M.Q. (2006). A WNT of things to come: evolution of Wnt signaling and polarity in cnidarians. Seminars in Cell & Developmental Biology 17, 157–167.

Liew, Y.J., Aranda, M., and Voolstra, C.R. (2016). Reefgenomics.org - a repository for marine genomics data. Database 2016.

Liew, Y.J., Howells, E.J., Wang, X., Michell, C.T., Burt, J.A., Idaghdour, Y., and Aranda, M. (2020). Intergenerational epigenetic inheritance in reef-building corals. Nature Climate Change 10, 254–259.

Love, M.I., Huber, W., and Anders, S. (2014). Moderated estimation of fold change and dispersion for RNA-Seq data with DESeq2. Genome Biology 15, 1–21.

Mackie, G. (1986). From aggregates to integrates: physiological aspects of modularity in colonial animals. Philosophical Transactions of the Royal Society of London. B, Biological Sciences 313, 175–196.

Malik, A., Einbinder, S., Martinez, S., Tchernov, D., Haviv, S., Almuly, R., Zaslansky, P., Polishchuk, I., Pokroy, B., and Stolarski, J. (2021). Molecular and skeletal fingerprints of scleractinian coral biomineralization: from the sea surface to mesophotic depths. Acta Biomaterialia 120, 263–276.

Marsh Jr, J.A. (1970). Primary productivity of reef-building calcareous red algae. Ecology 51, 255–263.

Martinez, S., Kolodny, Y., Shemesh, E., Scucchia, F., Nevo, R., Levin-Zaidman, S., Paltiel, Y., Keren, N., Tchernov, D., and Mass, T. (2020). Energy sources of the depth-generalist mixotrophic coral *Stylophora pistillata*. Frontiers in Marine Science 7, 988.

Mass, T., Brickner, I., Hendy, E., and Genin, A. (2011). Enduring physiological and reproductive benefits of enhanced flow for a stony coral. Limnology and Oceanography 56, 2176–2188.

Mass, T., Drake, Jeana L., Haramaty, L., Kim, J.D., Zelzion, E., Bhattacharya, D., and Falkowski, Paul G. (2013). Cloning and characterization of four novel coral acid-rich proteins that precipitate carbonates in vitro. Current Biology 23, 1126–1131.

Mass, T., Drake, J.L., Peters, E.C., Jiang, W., and Falkowski, P.G. (2014). Immunolocalization of skeletal matrix proteins in tissue and mineral of the coral *Stylophora pistillata*. Proceedings of the National Academy of Sciences 111, 12728–12733.

Mass, T., Einbinder, S., Brokovich, E., Shashar, N., Vago, R., Erez, J., and Dubinsky, Z. (2007). Photoacclimation of *Stylophora pistillata* to light extremes: metabolism and calcification. Marine Ecology Progress Series 334, 93–102.

Mass, T., Genin, A., Shavit, U., Grinstein, M., and Tchernov, D. (2010). Flow enhances photosynthesis in marine benthic autotrophs by increasing the efflux of oxygen from the organism to the water. Proceedings of the National Academy of Sciences 107, 2527–2531.

Mass, T., Putnam, H.M., Drake, J.L., Zelzion, E., Gates, R.D., Bhattacharya, D., and Falkowski, P.G. (2016). Temporal and spatial expression patterns of biomineralization proteins during early development in the stony coral *Pocillopora damicornis*. Proceedings of the Royal Society B: Biological Sciences 283, 20160322.

Matsuda, S.B., Huffmyer, A.S., Lenz, E.A., Davidson, J.M., Hancock, J.R., Przybylowski, A., Innis, T., Gates, R.D., and Barott, K.L. (2020). Coral bleaching susceptibility Is predictive of subsequent mortality within but not between coral species. Frontiers in Ecology and Evolution 8.

Meesters, E.H., and Bak, R.P. (1995). Age-related deterioration of a physiological function in the branching coral *Acropora palmata*. Marine Ecology Progress Series 121, 203–209.

Mummadisetti, M.P., Drake, J.L., and Falkowski, P.G. (2021). The spatial network of skeletal proteins in a stony coral. Journal of The Royal Society Interface 18, 20200859.

Nakamura, H., Asari, T., Ohizumi, Y., Kobayashi, J.i., Yamasu, T., and Murai, A. (1993). Isolation of zooxanthellatoxins, novel vasoconstrictive substances from the zooxanthella *Symbiodinium* sp. Toxicon 31, 371–376.

Palumbi, S.R., Barshis, D.J., Traylor-Knowles, N., and Bay, R.A. (2014). Mechanisms of reef coral resistance to future climate change. Science 344, 895–898.

Park, E., Hwang, D.-S., Lee, J.-S., Song, J.-I., Seo, T.-K., and Won, Y.-J. (2012). Estimation of divergence times in cnidarian evolution based on mitochondrial protein-coding genes and the fossil record. Molecular Phylogenetics and Evolution 62, 329–345.

Peled, Y., Drake, J.L., Malik, A., Almuly, R., Lalzar, M., Morgenstern, D., and Mass, T. (2020). Optimization of skeletal protein preparation for LC–MS/MS sequencing yields additional coral skeletal proteins in *Stylophora pistillata*. BMC Materials 2, 8.

Primor, N., and Zlotkin, E. (1975). On the ichthyotoxic and hemolytic action of the skin secretion of the flatfish *Pardachirus marmoratus* (Soleidae). Toxicon 13, 227–231.

Puverel, S., Tambutté, E., Pereira-Mouries, L., Zoccola, D., Allemand, D., and Tambutté, S. (2005). Soluble organic matrix of two Scleractinian corals: Partial and comparative analysis. Comparative Biochemistry and Physiology Part B 141, 480–487.

Rachamim, T., Morgenstern, D., Aharonovich, D., Brekhman, V., Lotan, T., and Sher, D. (2015). The dynamically evolving nematocyst content of an anthozoan, a scyphozoan, and a hydrozoan. Molecular Biology and Evolution 32, 740–753.

Ralph, P., Gademann, R., Larkum, A., and Kühl, M. (2002). Spatial heterogeneity in active chlorophyll fluorescence and PSII activity of coral tissues. Marine Biology 141, 639–646.

Ramos-Silva, P., Kaandorp, J., Huisman, L., Marie, B., Zanella-Cleon, I., Guichard, N., Miller, D.J., and Marin, F. (2013). The skeletal proteome of the coral *Acropora millepora:* The evolution of calcification by cooption and domain shuffling. Molecular Biology and Evolution.

Reinhardt, B., Broun, M., Blitz, I.L., and Bode, H.R. (2004). HyBMP5-8b, a BMP5-8 orthologue, acts during axial patterning and tentacle formation in *Hydra*. Developmental Biology 267, 43–59.

Rinkevich, B., and Loya, Y. (1983). Oriented translocation of energy in grafted reef corals. Coral Reefs 1, 243–247.

Ritchie, K.B. (2006). Regulation of microbial populations by coral surface mucus and mucus-associated bacteria. Marine Ecology Progress Series 322, 1–14.

Ritchie, R. (2008). Universal chlorophyll equations for estimating chlorophylls a, b, c, and d and total chlorophylls in natural assemblages of photosynthetic organisms using acetone, methanol, or ethanol solvents. Photosynthetica 46, 115–126.

Rocha, R.J., Silva, A.M., Fernandes, M.H.V., Cruz, I.C., Rosa, R., and Calado, R. (2014). Contrasting light spectra constrain the macro and microstructures of scleractinian corals. PLOS ONE 9.

Rueffler, C., Hermisson, J., and Wagner, G.P. (2012). Evolution of functional specialization and division of labor. Proceedings of the National Academy of Sciences 109, E326–E335.

Schiller, C., and Herndl, G.J. (1989). Evidence of enhanced microbial activity in the interstitial space of branched corals: possible implications for coral metabolism. Coral Reefs 7, 179–184.

Schmidt, C.A., Daly, N.L., and Wilson, D.T. (2019). Coral venom toxins. Frontiers in Ecology and Evolution 7, 320.

Schmidt-Roach, S., Miller, K.J., Lundgren, P., and Andreakis, N. (2014). With eyes wide open: a revision of species within and closely related to the *Pocillopora damicornis* species complex (Scleractinia; Pocilloporidae) using morphology and genetics. Zoological Journal of the Linnean Society 170, 1–33.

Sebens, K.P., Witting, J., and Helmuth, B. (1997). Effects of water flow and branch spacing on particle capture by the reef coral *Madracis mirabilis* (Duchassaing and Michelotti). Journal of Experimental Marine Biology and Ecology 211, 1–28.

Sergushichev, A.A. (2016). An algorithm for fast preranked gene set enrichment analysis using cumulative statistic calculation. BioRxiv, 060012.

Shenkar, N., Fine, M., and Loya, Y. (2005). Size matters: bleaching dynamics of the coral *Oculina patagonica*. Marine Ecology Progress Series 294, 181–188.

Siegfried, E., and Perrimon, N. (1994). *Drosophila* wingless: a paradigm for the function and mechanism of Wnt signaling. Bioessays 16, 395–404.

Steinmetz, P.R. (2019). A non-bilaterian perspective on the development and evolution of animal digestive systems. Cell and Tissue Research 377, 321–339.

Stimson, J. (1978). Mode and timing of reproduction in some common hermatypic corals of Hawaii and Enewetak. Marine Biology 48, 173–184.

Subramanian, A., Tamayo, P., Mootha, V.K., Mukherjee, S., Ebert, B.L., Gillette, M.A., Paulovich, A., Pomeroy, S.L., Golub, T.R., and Lander, E.S. (2005). Gene set enrichment analysis: a knowledge-based approach for interpreting genome-wide expression profiles. Proceedings of the National Academy of Sciences 102, 15545–15550.

Sunagawa, S., DeSalvo, M.K., Voolstra, C.R., Reyes-Bermudez, A., and Medina, M. (2009). Identification and gene expression analysis of a taxonomically restricted cysteine-rich protein family in reef-building corals. PLOS ONE 4, e4865.

Surekha, K.L., Khade, S., Trimbake, D., Patwardhan, R., Nadimpalli, S.K., and Ghaskadbi, S. (2020). Differential expression of BMP inhibitors gremlin and noggin in *Hydra* suggests distinct roles during budding and patterning of tentacles. Developmental Dynamics.

Takeuchi, T., Yamada, L., Shinzato, C., Sawada, H., and Satoh, N. (2016). Stepwise evolution of coral biomineralization revealed with genome-wide proteomics and transcriptomics. PLOS ONE 11, e0156424.

RStudio Team. (2019). “RStudio: Integrated Development for R”. (Boston, MA: RStudio, Inc.).

Voolstra, C.R., Li, Y., Liew, Y.J., Baumgarten, S., Zoccola, D., Flot, J.-F., Tambutté, S., Allemand, D., and Aranda, M. (2017). Comparative analysis of the genomes of *Stylophora pistillata* and *Acropora digitifera* provides evidence for extensive differences between species of corals. Scientific Reports 7, 17583.

Walden, P.D., and Cowan, N.J. (1993). A novel 205-kilodalton testis-specific serine/threonine protein kinase associated with microtubules of the spermatid manchette. Molecular and Cellular Biology 13, 7625–7635.

Wallace, C.C. (1978). The coral genus *Acropora* (scleractinia: Astrocoaniina: Acroporidae) in the central and southern Great Barrier Reef province. Memoires of the Queensland Museum 18, 273–319.

Wallace, C.C. (1985). Reproduction, recruitment and fragmentation in nine sympatric species of the coral genus *Acropora*. Marine Biology 88, 217–233.

Wangpraseurt, D., Polerecky, L., Larkum, A.W., Ralph, P.J., Nielsen, D.A., Pernice, M., and Kühl, M. (2014). The in situ light microenvironment of corals. Limnology and Oceanography 59, 917–926.

Winkler, R., Antonius, A., and Abigail Renegar, D. (2004). The skeleton eroding band disease on coral reefs of Aqaba, Red Sea. Marine Ecology 25, 129–144.

Xu, X., Weinstein, M., Li, C., and Deng, C.-X. (1999). Fibroblast growth factor receptors (FGFRs) and their roles in limb development. Cell and Tissue Research 296, 33–43.

Yahel, R., Yahel, G., Berman, T., Jaffe, J.S., and Genin, A. (2005). Diel pattern with abrupt crepuscular changes of zooplankton over a coral reef. Limnology and Oceanography 50, 930–944.

Yosef, O., Popovits, Y., Malik, A., Ofek-Lalzer, M., Mass, T., and Sher, D. (2020). A tentacle for every occasion: comparing the hunting tentacles and sweeper tentacles, used for territorial competition, in the coral *Galaxea fascicularis*. BMC Genomics 21, 1–16.

Young, M.D., Wakefield, M.J., Smyth, G.K., and Oshlack, A. (2012). goseq: Gene Ontology testing for RNA-seq datasets. R Bioconductor 8, 1–25.

Yu, G., Lam, T.T.-Y., Zhu, H., and Guan, Y. (2018). Two methods for mapping and visualizing associated data on phylogeny using ggtree. Molecular Biology and Evolution 35, 3041–3043.

Zaquin, T., Malik, A., Drake, J.L., Putnam, H.M., and Mass, T. (2021). Evolution of protein-mediated biomineralization in scleractinian corals. Frontiers in Genetics 12, 52.

Zoccola, D., Ganot, P., Bertucci, A., Caminiti-Segonds, N., Techer, N., Voolstra, C.R., Aranda, M., Tambutté, E., Allemand, D., and Casey, J.R. (2015). Bicarbonate transporters in corals point towards a key step in the evolution of cnidarian calcification. Scientific Reports 5.

Zoccola, D., Innocenti, A., Bertucci, A., Tambutté, E., Supuran, C.T., and Tambutté, S. (2016). Coral carbonic anhydrases: regulation by ocean acidification. Marine Drugs 14, 109.

