## Supplementary Information for "Physiological and transcriptomic variability indicative of differences in key functions within a single coral colony"

Supplementary Information: Extended Methods, Table Legends, Figures, and Figure Legends for: Physiological and transcriptomic variability indicative of differences in key functions within a single coral colony

Jeana L. Drake[1+], Assaf Malik[1+], Yotam Popovits[1+], Oshra Yosef[1], Eli Shemesh[1], Jarosław Stolarski[2], Dan Tchernov[1,3], Daniel Sher[1*] and Tali Mass[1,3*]

[1]Department of Marine Biology, University of Haifa, 199 Aba Koushy Avenue, Haifa, Israel, 3498838
[2]Institute of Paleobiology, Polish Academy of Sciences, Twarda 51/55, Warsaw, Poland, 00-818
[3]Morris Kahn Marine Research Station, The Leon H. Charney School of Marine Sciences, University of Haifa, Sdot Yam, Israel

[+]These authors contributed equally
[*]Corresponding authors

### 1 Supplementary Tables

The data underlying this article are available in the article, in its online Supplementary Material, and on GitHub in the following repository: https://github.com/jeanadrake/Single_coral_colony_transcriptomes_physiology.

Supplementary Table 1. Dissolved oxygen concentration and total SYBR-stained or chlorophyll auto-fluorescence-based microbial cell counts and measured inside and outside five *S. pistillata* colonies in situ.

Supplementary Table 2. Statistical analysis of environmental parameters measured between branches and at colony peripheries, and physiological parameters measured at two tips, a junction, and a base segment on each of five branches distributed throughout the *S. pistillata* colony.

Supplementary Table 3. Location information for branches used for transcriptional analysis. DESeq2 analysis used the additive model output so that the top-most branch or 9$^{th}$ branch, which points upward and thus does not have a defined cardinal direction, was not included.

Supplementary Table 4. DE gene count statistics. The counts of over-expressed/under-expressed/unchanged/ NA genes using DESeq2 additive and interaction models. Also included are read count and filtered genes statistics. Both host *Stylophora pistillata* and photosymbiont (reads mapped to the *Symbiodinium microadriaticum* genome, NCBI GCA_001939145.1) data are shown.

Supplementary Table 5. *S. pistillata* additive model DE. DESeq2 additive model statistics of DE and gene annotations.

Supplementary Table 6. *S. pistillata* interaction model DE. DESeq2 interaction model statistics of DE and gene annotations.

Supplementary Table 7. Photosymbiont additive model DE. DESeq2 additive model statistics of DE and gene annotations.

Supplementary Table 8. Host and photosymbiont factor effect Adonis (non-parametric multivariate analysis of variance, known as NPMANOVA or PERMANOVA). The Adonis test, implemented in Vegan, was used to test the effect of factors branch axis, ring height, and cardinal direction, based on NMDS two-dimensional sample clustering.

Supplementary Table 9. *S. pistillata* vs . *Acropora* spp. (Hemond et al.) DE tip vs base statistics for putative orthologous genes found using reciprocal BLAST.

Supplementary Table 10. *S. pistillata* KEGG and GO (Trinotate) enrichment based on the DESeq2 additive-model. Enrichment analysis uses the GOSeq Wallenius (Wall) method.

Supplementary Table 11. *S. pistillata* KEGG and GO (Trinotate) enrichment based on the DESeq2 interaction-model. Enrichment analysis uses the GOSeq Wallenius (Wall) method.

Supplementary Table 12. - Photosymbiont GO enrichment (Uniprot database) of DE genes based on the DESeq2 additive-model. Enrichment analysis uses the Gene Set Enrichment Analysis (GSEA) method.

Supplementary Table 13. Physiological parameters measured along five branches from the same single colony of *S. pistillata* from which differential gene expression was also examined.

Supplementary Table 14. DE of putative toxins sequenced in this study obtained by reciprocal blast against known toxins from the literature.

Supplementary Table 15. DE of orthologous putative toxins between *S. pistillata* and *Acropora* spp (Hemond et al.). Most putative toxins from *S. pistillata* do not have orthologs in the *Acropora* spp. dataset.

Supplementary Table 16. DE of known biomineralization-related genes. 60 genes code for proteins recently sequenced from *S. pistillata* skeleton (Peled et al., 2020). CARPs 1, 2, and 3 (Mass et al., 2013) and a putative bicarbonate transporter (Zoccola et al., 2015) were appended to the list. Not all genes associated with known coral skeletal proteins were sequenced here.

Supplementary Table 17. DE of orthologous biomineralization-related genes between *S. pistillata* and *Acropora* spp (Hemond et al.). Just under half of such genes from *S. pistillata* have orthologs in the *Acropora* spp. dataset.

### 2 Supplementary Figures

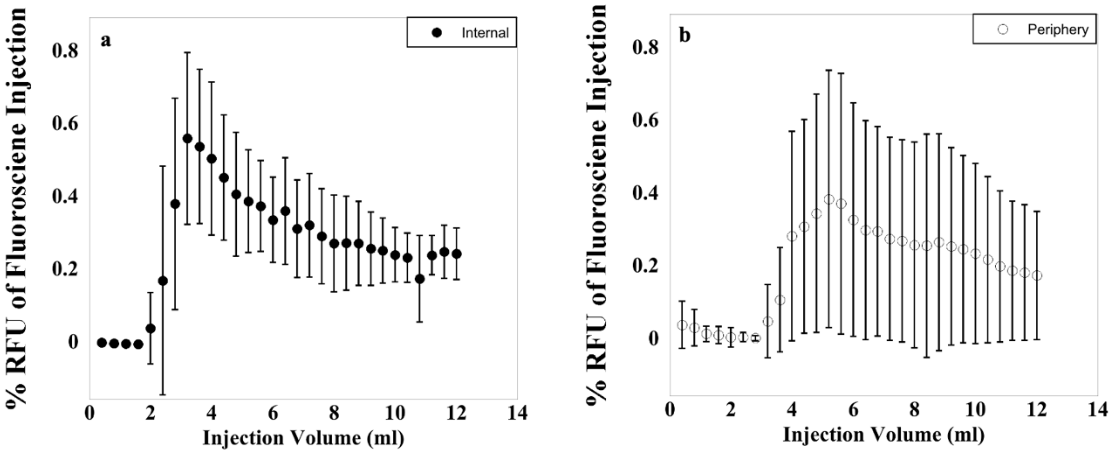


Supplementary Figure 1. Percent fluorescence, relative to the injected fluoresceine dye aliquot's fluorescence, of various volumes of water extracted by syringe either on the opposite side of the branch from the dye injection (a) or at colony peripheries (b). Fluorescence higher than the background water was not observed for extraction volumes of 1.6 ml or smaller.

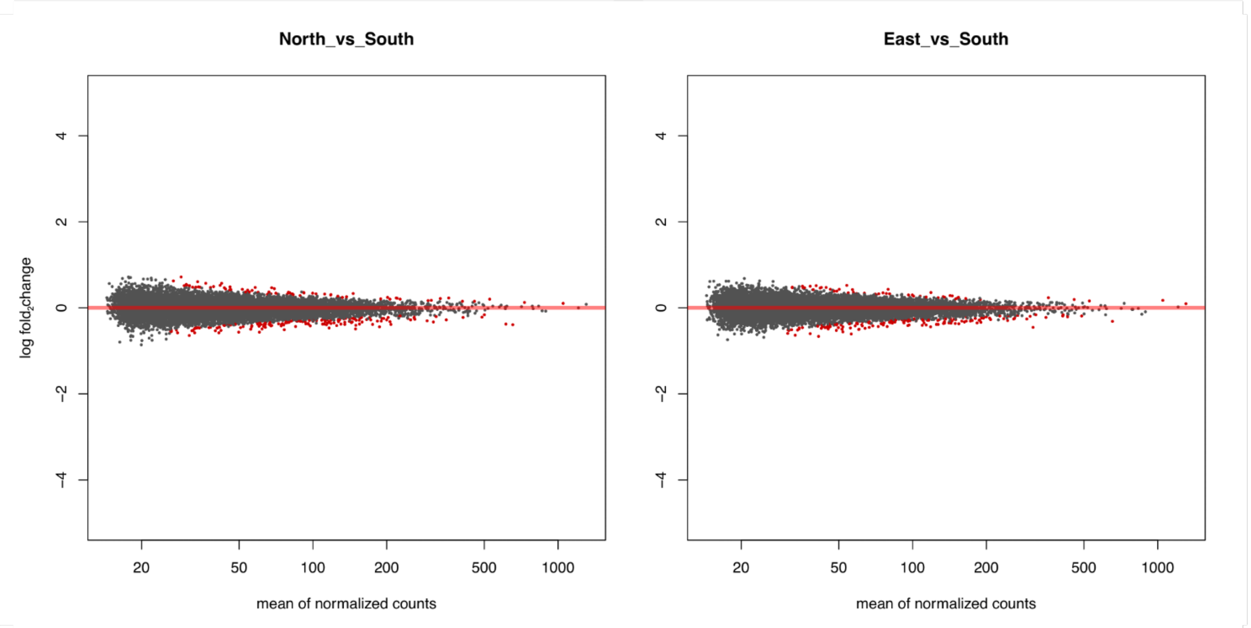


Supplementary Figure 2. Photosymbiont MA plots. The plots show the genes' mean normalized counts vs. $log_2$fold change, where red dots represent significant DE genes, for North-South and East-South comparisons.

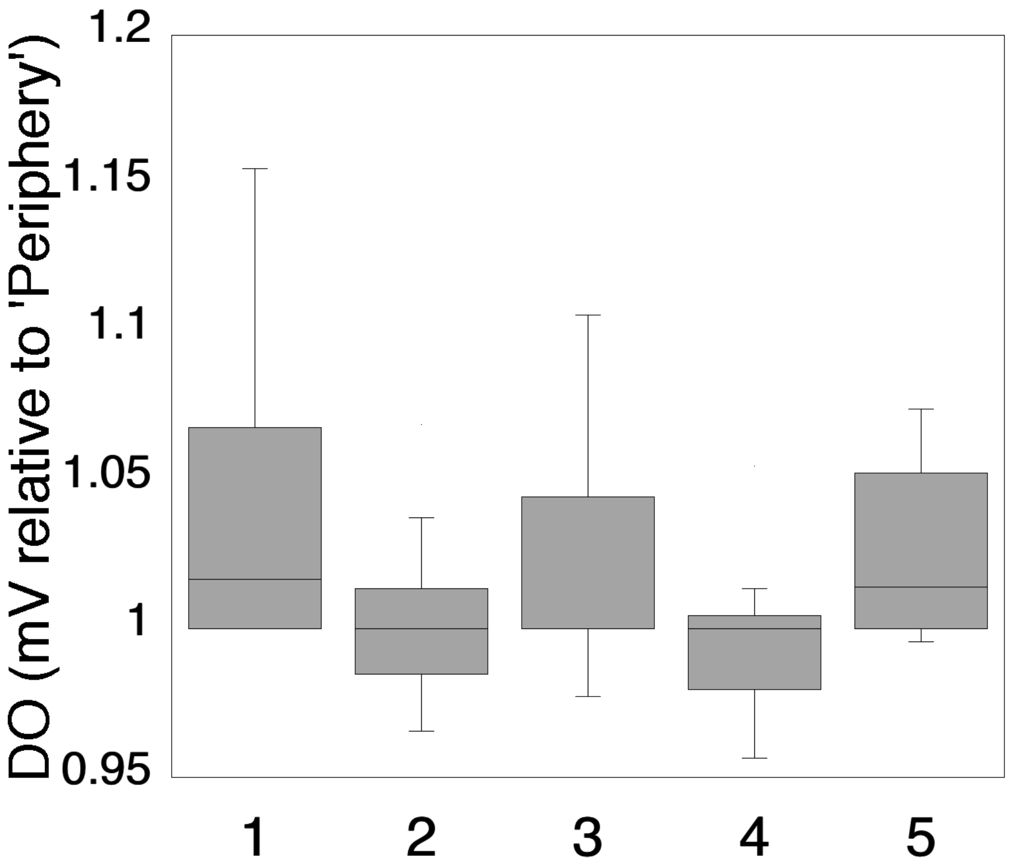


Supplementary Figure 3. Normalized mV measurements as proxies of DO measured along transects from inside to just outside five coral colonies growing in the same nursery as the colony examined at high spatial resolution for transcriptional, physiological, and morphological differences. Measurements for each colony were standardized to the ‘Periphery’ measurement of each transect for statistical analysis of DO differences by general location within a colony.

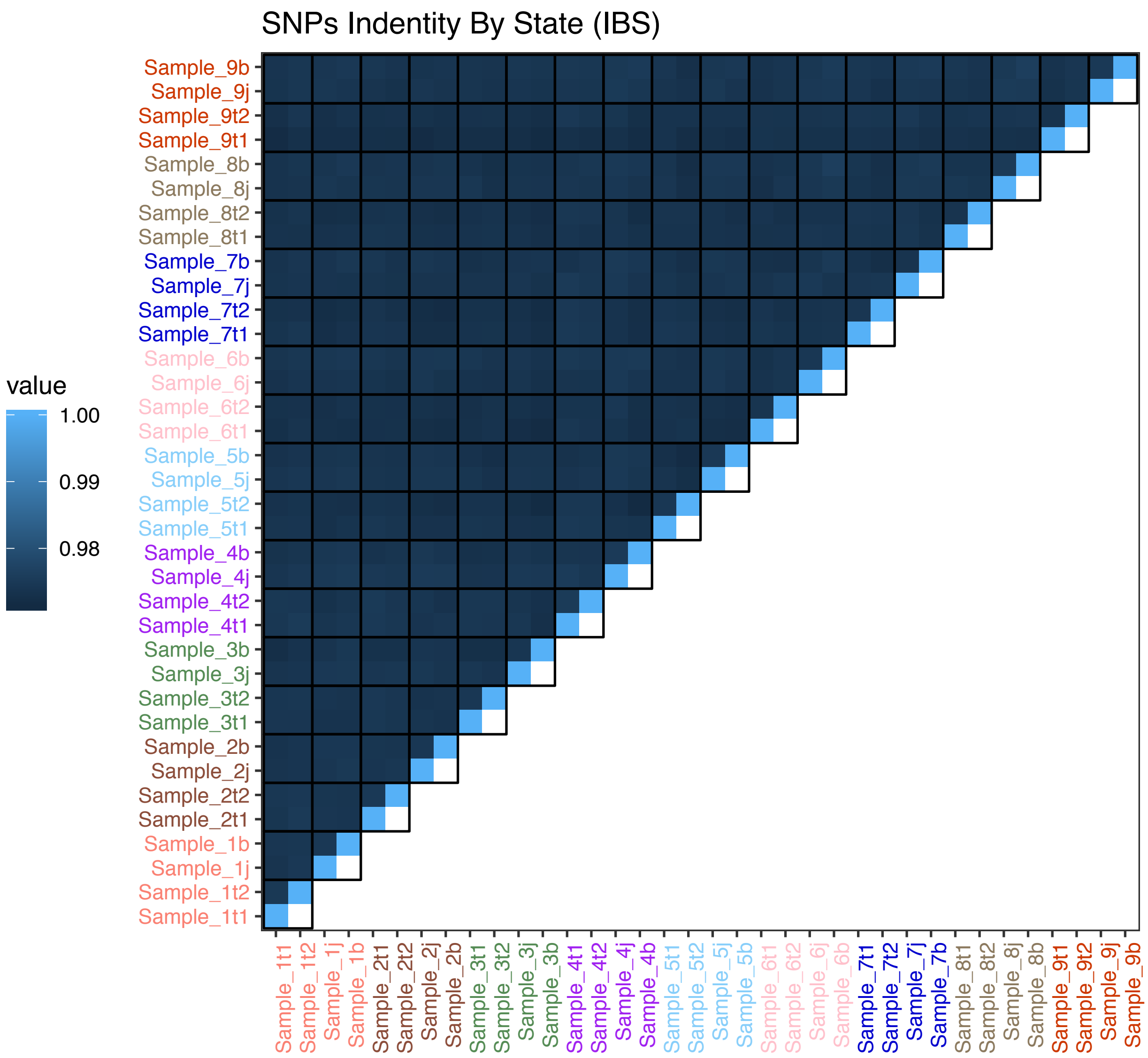


Supplementary Figure 4. Single nucleotide polymorphism (SNP) identity by state. SNP identity by state heatmap showing the level of SNP identity between samples (0-1.0 indicates 0-100% identity), based on the Broad Institute's GATK.

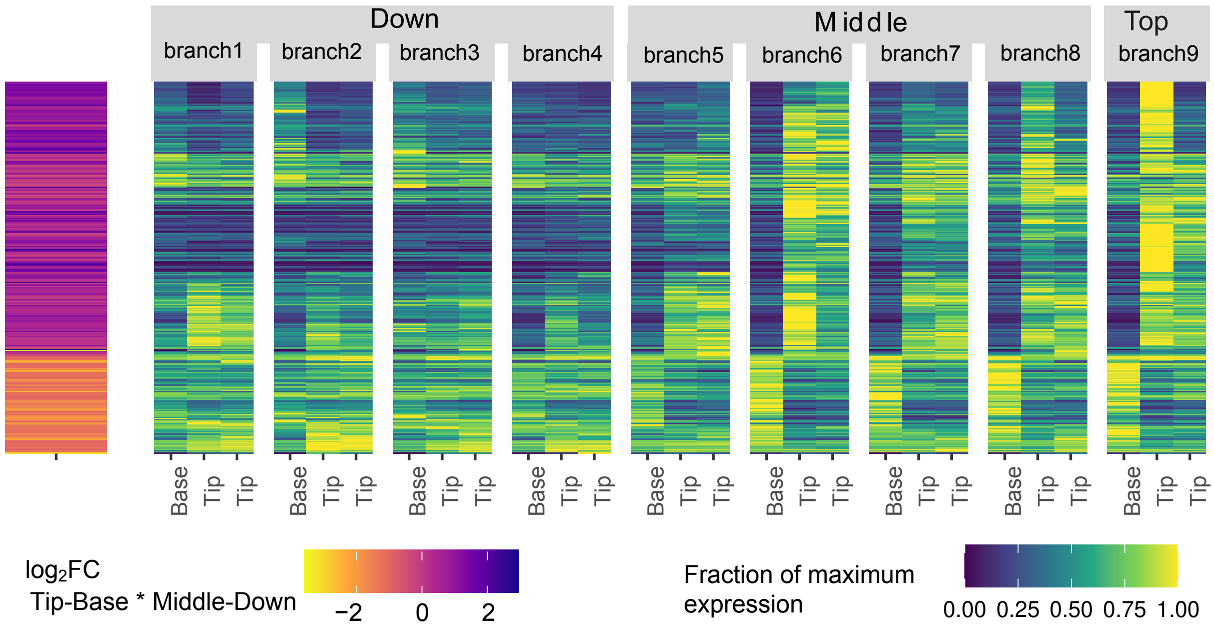


Supplementary Figure 5. Interaction heatmap of host genes. The heatmap on the left shows the relative expression values of a subset of *S. pistillata* genes that demonstrate a significant interaction between base-tip (position factor) versus middle-bottom (ring height factor). The heatmap on the right represents the $\log_2$FC of the interactions. For clarity, only interactions for tips and bases are shown.

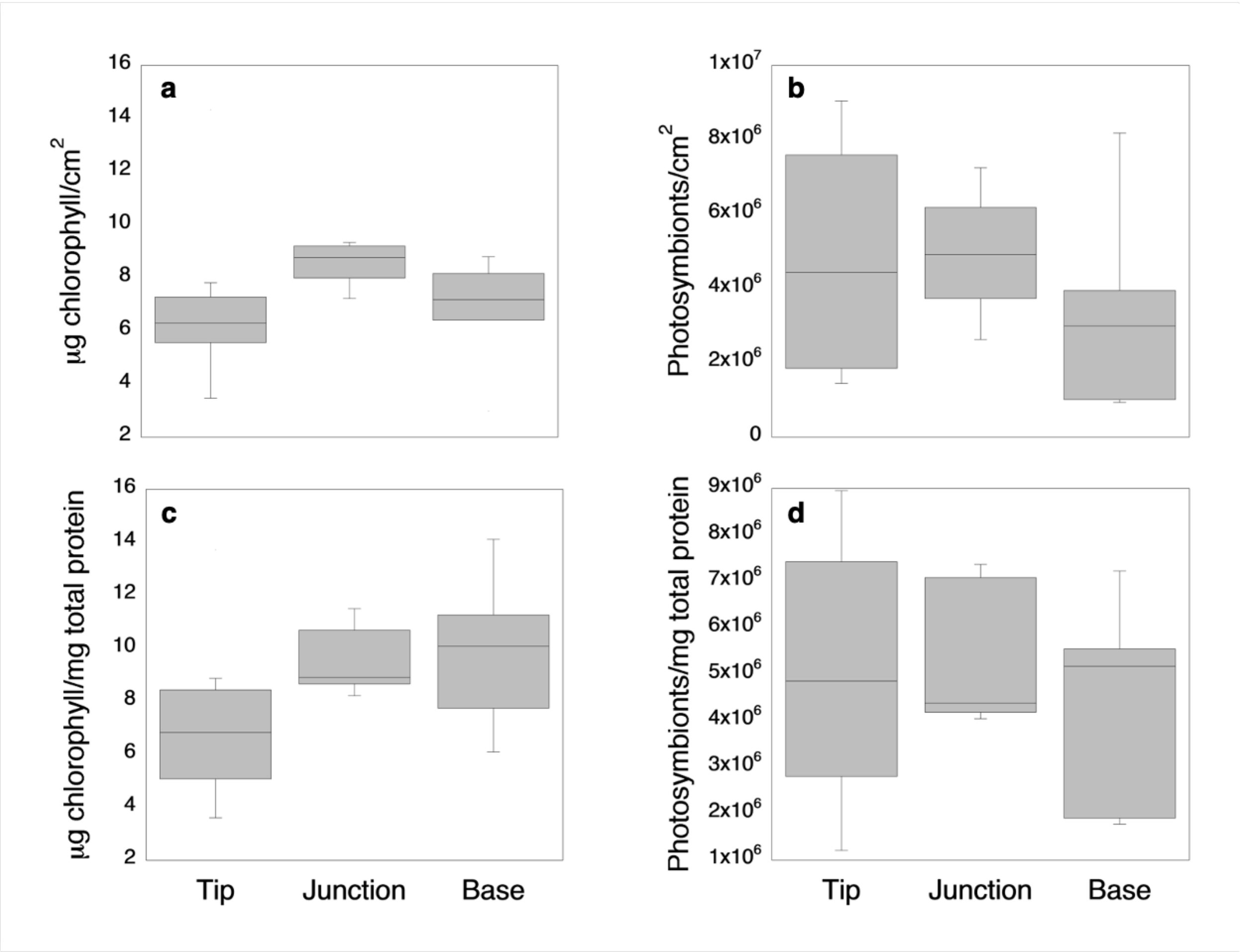


Supplementary Figure 6. Select measures of photophysiology were not significantly different between branch locations across the single *S. pistillata* colony measured here, as chlorophyll content (a, c) and photosymbiont abundance (b, d) by either surface area (a, b) or total protein (c, d).

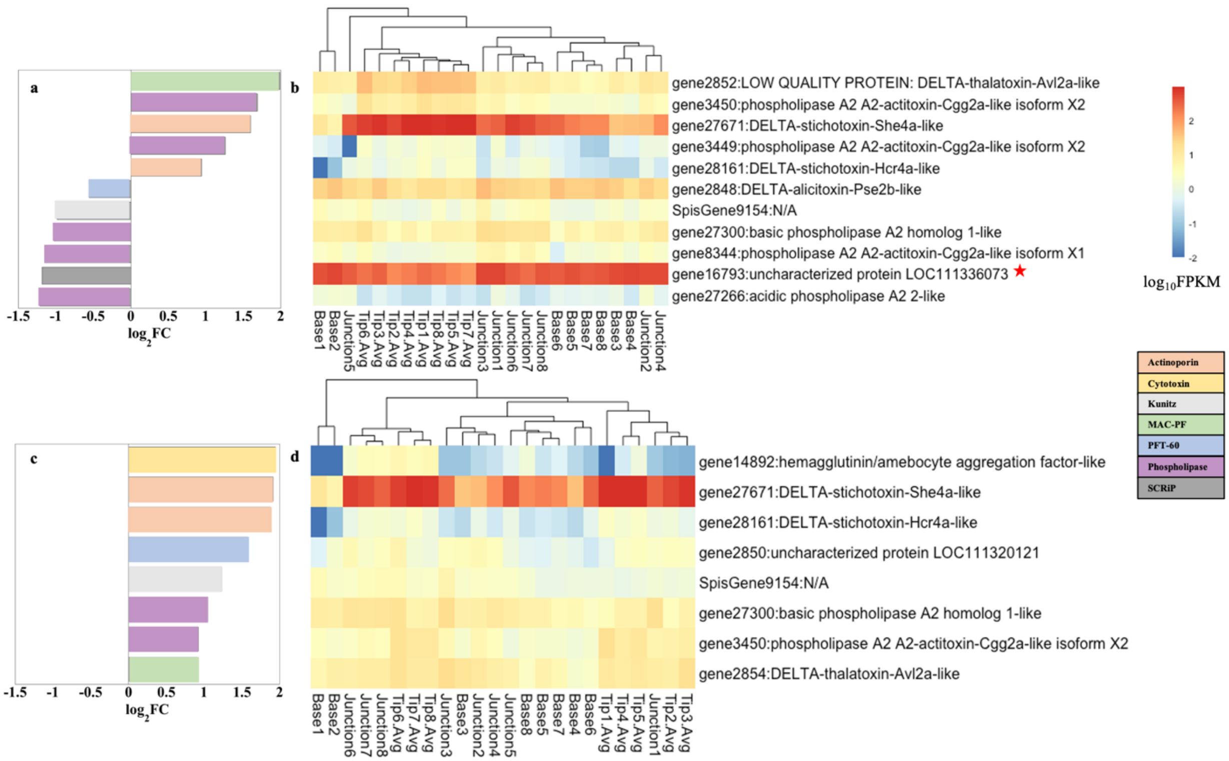


Supplementary Figure 7. Gene expression patterns of putative toxins as $\log_2$FC (a, c), and $\log_{10}$FPKM (b, d) between branch tips and junctions (a, b) and junctions and bases (c, d). Only significantly DE toxin genes are shown ($p<0.05$). An uncharacterized gene, for which reciprocal BLAST hits suggest it is a SCRiP (Supplementary Table 14), is noted with a red star (b).

a

| Query | Best Blast Hit | E-value | % Identity | BLAST Server |
|---|---|---|---|---|
| Stylophora_pistillata_9725 | XP_022777760.1 | 8 e-122 | 87.4 | NCBI |
| gene27671 | XP_022777760.1 | 0 | 100 | NCBI |
| XP_022777760.1 | Stylophora_pistillata_9725 | 7.4 e-126 | 87.4 | Comparative Reef Genomics |
| gene27671 | Stylophora_pistillata_9725 | 7.4 e-126 | 87 | Comparative Reef Genomics |

b

```
gene27671_rna35563              MEMKMCALLSFYILAVCFMGYCVSGAHIEFKEDFKEMGNVKRDMGAHLETSNPIGETENT    60
XP_022777760.1                  MEMKMCALLSFYILAVCFMGYCVSGAHIEFKEDFKEMGNVKRDMGAHLETSNPIGETENT    60
Stylophora_pistillata_9725      --------------------------------------------------------MENT    4
                                                                                         ***

gene27671_rna35563              ERELARQHQQKKRVNHATVIDAASLTKDVLQTLLDALGSIERKIAIGIANETPYEWRALN    120
XP_022777760.1                  ERELARQHQQKKRVNHATVIDAASLTKDVLQTLLDALGSIERKIAIGIANETPYEWRALN    120
Stylophora_pistillata_9725      ERELAKKHNQNKRVDPGTVIAVASLTKELLQTLLDALGSIERKIAIGIANETPYEWRALN    64
                                *****::*:*:***: .*** .*****::*******************************

gene27671_rna35563              TYFRSGTSDDLLPPFVKKDQAPLYTARKTNGPVATGCVAVIAYYMPAVQMTVGVMFSVPF    180
XP_022777760.1                  TYFRSGTSDDLLPPFVKKDQAPLYTARKTNGPVATGCVAVIAYYMPAVQMTVGVMFSVPF    180
Stylophora_pistillata_9725      TYFRSGTSDDLLPRFVEKDQAALYTARKTNGPVATGCVAVIAYYMPTVQMTVGVMFSVPF    124
                                ************* **:**** ************************:*************

gene27671_rna35563              DQNFYRNWWNARVYPGEVRASQRMYEDMYYGNPFRGDNGWHKKDIGHGFHMDGSMTSPGK    240
XP_022777760.1                  DQNFYRNWWNARVYPGEVRASQRMYEDMYYGNPFRGDNGWHKKDIGHGFHMDGSMTSPGK    240
Stylophora_pistillata_9725      DYNFYSNWWDARVYHGELRASQQMYEDMYYGNPFRGDNGWHKKDIGYGFHVDGSMTSAGK    184
                                * *** ***:**** **:****:***********************:***:****** **

gene27671_rna35563              SVMELHITSN      250
XP_022777760.1                  SVMELHITSN      250
Stylophora_pistillata_9725      SVMELHILNK      194
                                ******* .:
```

Supplementary Figure 8. Reciprocal protein blast information (a) and sequence alignment (b) for the D-Pocilopotoxin-Spi1 in (Ben-Ari et al., 2018) (Stylophora_pistillata_9725 from (Bhattacharya et al., 2016)) and the gene27671 (rna35563) annotated here as DELTA-stichotoxin-She4a-like (NCBI accession number XP_022777760.1).

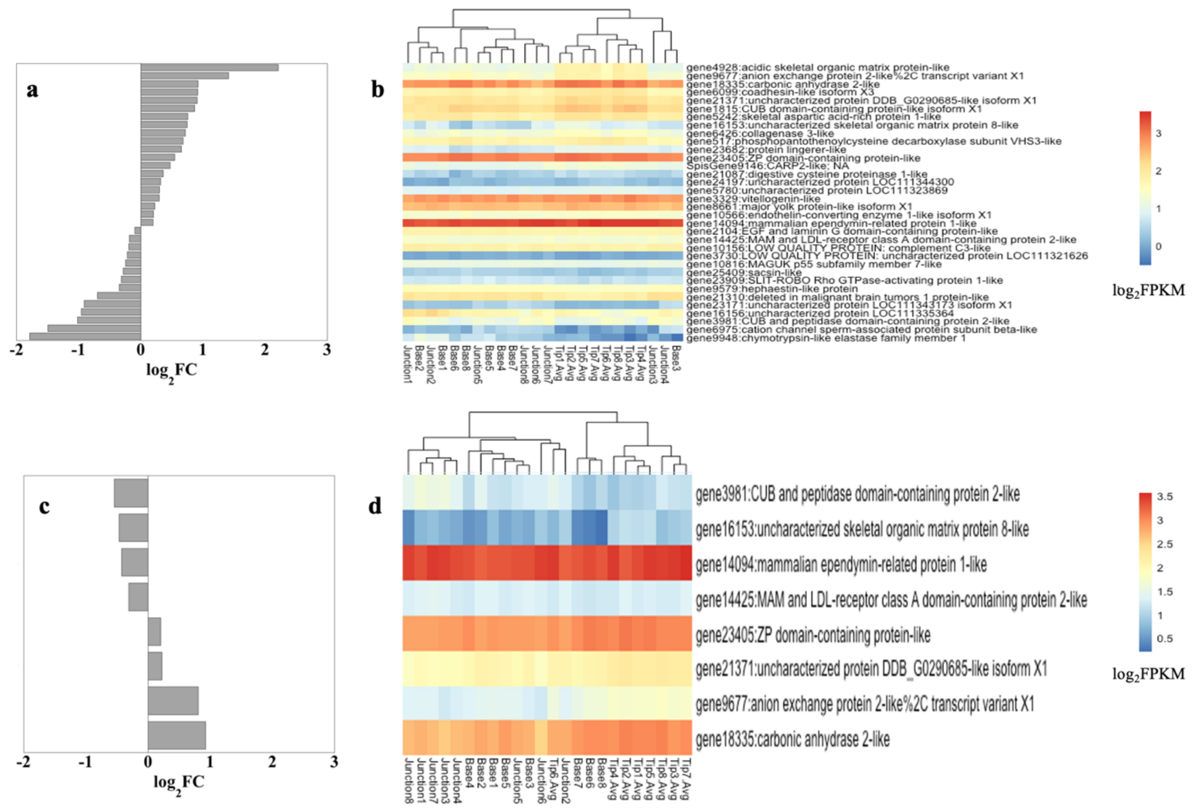


Supplementary Figure 9. Gene expression patterns of known biomineralization-related genes as $log_2FC$ (a, c), and $log_{10}FPKM$ (b, d) between branch tips and junctions (a, b) and junctions and bases (c, d). Only significantly DE biomineralization genes are shown ($p<0.05$).

### 3 Data Availability Statement

The data underlying this article are available in the article, in its online Supplementary Material, and on GitHub in the following repository: https://github.com/jeanadrake/Single_coral_colony_transcriptomes_physiology.

### 4 References


Ben-Ari, H., Paz, M., and Sher, D. (2018). The chemical armament of reef-building corals: inter- and intra-specific variation and the identification of an unusual actinoporin in *Stylophora pistilata*. *Scientific Reports* 8**,** 1-13.

Bhattacharya, D., Agrawal, S., Aranda, M., Baumgarten, S., Belcaid, M., Drake, J.L., Erwin, D., Foret, S., Gates, R.D., Gruber, D.F., Kamel, B., Lesser, M.P., Levy, O., Liew, Y.J., MacManes, M., Mass, T., Medina, M., Mehr, S., Meyer, E., Price, D.C., Putnam, H.M., Qiu, H., Shinzato, C., Shoguchi, E., Stokes, A.J., Tambutté, S., Tchernov, D., Voolstra, C.R., Wagner, N., Walker, C.W., Weber, A.P., Weis, V.M., Zelzion, E., Zoccola, D., and Falkowski, P. (2016). Comparative genomics explains the evolutionary success of reef-forming corals. *eLife* 5**,** e13288.

Hemond, E.M., Kaluziak, S.T., and Vollmer, S.V. (2014). The genetics of colony form and function in Caribbean *Acropora* corals. *BMC Genomics* 15**,** 1133.

Mass, T., Drake, Jeana L., Haramaty, L., Kim, J.D., Zelzion, E., Bhattacharya, D., and Falkowski, Paul G. (2013). Cloning and characterization of four novel coral acid-rich proteins that precipitate carbonates in vitro. *Current Biology* 23**,** 1126-1131.

Peled, Y., Drake, J.L., Malik, A., Almuly, R., Lalzar, M., Morgenstern, D., and Mass, T. (2020). Optimization of skeletal protein preparation for LC–MS/MS sequencing yields additional coral skeletal proteins in *Stylophora pistillata*. *BMC Materials* 2**,** 8.

Zoccola, D., Ganot, P., Bertucci, A., Caminiti-Segonds, N., Techer, N., Voolstra, C.R., Aranda, M., Tambutté, E., Allemand, D., and Casey, J.R. (2015). Bicarbonate transporters in corals point towards a key step in the evolution of cnidarian calcification. *Scientific Reports* 5.
